# Altered Physiology and Ensemble Recruitment of Dentate Gyrus Semilunar Granule Cells in a Mouse Model of Epilepsy

**DOI:** 10.64898/2026.01.24.701531

**Authors:** Laura Dovek, Andrew Huang, Vijayalakshmi Santhakumar

## Abstract

The dentate gyrus is a major locus for structural and synaptic reorganization in temporal lobe epilepsy. While physiological changes during epileptogensis are well characterized in the principal dentate projection neuron, granule cells (GCs), epilepsy-related changes in semilunar granule cells (SGCs), a distinct subset of dentate projection neurons, remain unknown. Using a mouse pilocarpine model of epilepsy, we show that, unlike GCs, SGCs exhibit an increase in intrinsic excitability 1 week after status epilepticus (SE). GCs, not SGCs. have a lower threshold for action potential firing in mice 1-month post-SE. Both GCs and SGCs display increased frequency of spontaneous excitatory postsynaptic currents (EPSCs) 1-week and 1-month after SE. Uniquely, SGCs received more frequent spontaneous inhibitory postsynaptic currents (sIPSCs) and exhibited smaller afferent-evoked IPSCs early after SE. Additionally, evoked EPSC amplitude in SGCs was reduced 1-month post-SE. Behaviorally, mice 1-month post-SE showed impairments in their ability to use spatial search strategies in a Barnes maze paradigm. Post-SE TRAP2::tdT mice showed reduced activity dependent neuronal ensemble labeling, with fewer tdT-labeled neurons in both task-naïve and trained conditions and reduced *c-Fos* co-expression following task re-acquisition compared to controls. Notably, the proportion of SGCs within labeled ensembles was reduced in task-naïve epileptic mice but not in trained animals. Collectively, our findings identify selective changes in SGC intrinsic excitability during epileptogenesis that could contribute to enhanced network excitability. The cell-specific alterations in SGC circuit connectivity during epileptogenesis, alongside the apparent reduction in neuronal recruitment to behavioral ensembles likely contribute to spatial navigation deficits observed in epilepsy.

## Introduction

Temporal lobe epilepsy (TLE) is the most prevalent focal epilepsy, with approximately 100,000 new cases diagnosed annually, yet a third of these cases are refractory to current pharmacological treatments (Daoud et al., 2004; Laxer et al., 2014). The ensuing cognitive and psychological comorbidities significantly impact the quality of life of TLE patients. The dentate gyrus (DG) is central to memory processing (Senzai, 2019; Hainmueller and Bartos, 2020; GoodSmith et al., 2022) and the primary site for circuit level changes in epilepsy (Scharfman and Pedley, 2007; Dengler and Coulter, 2016; Scharfman, 2019). The DG receives extensive inputs from the entorhinal cortex and processes them into sparse, temporally precise patterns of cellular activity which are then transmitted downstream through the hippocampal trisynaptic circuit for memory consolidation. This multi-stage processing of inputs is considered critical for episodic memory formation, which involves the integration of multiple sensory modalities (Liu et al., 2012; Liu et al., 2014; Guan et al., 2016; Hainmueller and Bartos, 2018, 2020). Following epileptogenic insults, the DG experiences cell death, aberrant axonal sprouting, and circuit reorganization that alter network excitability and impair memory processing (Houser and Esclapez, 1996; Buckmaster and Dudek, 1997; Buckmaster et al., 2002; Zhang et al., 2009; Dengler and Coulter, 2016; Houser, 2024). Considering the impact of epileptogenesis on memory function, there is a pressing need to understand how dentate circuit function is regulated in both healthy and epileptic states.

Although the DG circuit and granule cells (GCs), the canonical projection neurons, have been examined extensively in epileptogenesis, recent functional analyses of a distinct projection neuron subtype suggest that the DG circuit may be more complex than previously proposed (Williams et al., 2007; Gupta et al., 2020; Afrasiabi et al., 2022; Dovek et al., 2025a). Semilunar granule cells (SGCs) are a sparse GC subtype distinguished by wide dendritic arbors, a “semilunar” shaped somata, and persistent firing properties. SGCs also display several differences in physiology, connectivity and plasticity compared to GCs, suggesting a unique role in circuit function (Williams et al., 2007; Larimer and Strowbridge, 2010; Gupta et al., 2020; Afrasiabi et al., 2022; Dovek et al., 2025a). Importantly, SGCs receive more frequent spontaneous excitatory and inhibitory postsynaptic events than GCs (Gupta et al., 2012; Gupta et al., 2020; Afrasiabi et al., 2022; Dovek et al., 2025a) and are preferentially recruited as part of neuronal ensembles labeled based on c-*Fos* expression (Erwin et al., 2020; Dovek et al., 2025b; Dovek et al., 2025a). SGCs also receive stronger associational glutamatergic drive from the hilus and medial entorhinal cortex (Williams et al., 2007; Dovek et al., 2025a), and exhibit sustained depolarizations, allowing them to engage hilar neurons during up-states (Williams et al., 2007). Furthermore, SGCs have been proposed to contribute to the refinement of neuronal ensembles through polysynaptic feedback inhibition (Larimer and Strowbridge, 2010; Walker et al., 2010; Erwin et al., 2020), with recent studies indicating that greater intrinsic excitability and shared synaptic drive contribute to recruitment of SGCs to active neuronal ensembles (Dovek et al., 2025b).

Although SGCs and GCs have been shown to diverge in how their tonic and synaptic inhibitory currents change over time after brain injury (Gupta et al., 2012; Gupta et al., 2022), SGC function and plasticity in epilepsy have not been examined. Prior studies have revealed alterations in synaptic inputs to GCs after status epilepticus (SE) and identified that the changes evolve over time with disease progression (Buckmaster and Dudek, 1997; Thind et al., 2008; Zhang et al., 2012; Althaus et al., 2019). Specifically, sprouting of GC mossy fiber axons into the inner molecular layer (IML), has been attributed to the loss of glutamatergic mossy cells (Houser et al., 1990; Cronin et al., 1992; Buckmaster et al., 2002; Jiao and Nadler, 2007) and has been implicated in increased excitatory drive onto GCs, heightened network activity and seizure generation (Buckmaster et al., 2002; Thind et al., 2008). These structural and functional changes to the DG circuit contribute to deficits in dentate dependent memory tasks performance (Gröticke et al., 2007; Bui et al., 2018; Kahn et al., 2019). However, it remains unclear whether these network level changes modify SGC function and whether epilepsy impacts recruitment of GCs versus SGCs to neuronal ensembles.

This study was conducted to determine whether the intrinsic physiology and synaptic inputs to SGCs and recruitment of SGCs to dentate memory-related neuronal ensembles are altered in experimental epilepsy.

## Methods

### Animals

All experiments were conducted under IACUC protocols approved by the University of California at Riverside and conformed with ARRIVE guidelines. Male and female wild type C57BL/6J (WT) mice were used for all electrophysiology experiments. c-*Fos*-cre mice (TRAP2: Fostm2.1^(icre/ERT2)Luo/J^); (Jackson Laboratories #030323) were back-crossed with C57BL6/N and were bred with floxed reporter line tdT-Ai14 mice (B6;129S6-Gt(ROSA)^26Sortm14(CAG-tdTomato)Hze/J^) (Jackson Laboratories # 007908) to create TRAP2::tdT mice. Mice were housed with littermates (up to 5 mice per cage) in a 12/12h light/dark cycle. WT mice, and a combination of TRAP2::tdT as well as TRAP2::ChR2-eYFP, collectively referred to as TRAP2::reporter mice, were used for EEG experiments to evaluate development of spontaneous recurrent seizures after status epilepticus (SE) induction. Mice used in behavioral studies were separated into individual cages one night prior to behavioral testing as pre-habituation. Food and water were provided ad libitum.

### Pilocarpine Status Epilepticus

Young TRAP2::reporter and WT mice postnatal days 28-35 mice were treated with scopolamine methyl nitrate (1mg/kg s.c.) 30 min before pilocarpine injection. SE was induced by injection of an initial dose of pilocarpine at 280mg/kg for TRAP2::reporter mice and 250mg/kg for *WT* mice followed by up to 3 additional half doses to induce stage 4 or 5 seizures according to the Racine scale (Racine, 1972). Dosage used to induce SE differed between the two mouse lines which is consistent with previously reported strain differences in pilocarpine sensitivity (Müller et al., 2009). Diazepam (10mg/kg, i.p.) was administered 60 minutes after the first stage 4 or 5 seizures. Only animals that had 3 or more stage 4 or 5 seizures during the pilocarpine administration were used in experiments (Proddutur et al., 2023). Control animals were littermates which received saline (i.p.) after scopolamine treatment followed by diazepam after 1.5 hours.

### Electroencephalograms (EEG)

A cohort of *WT* and TRAP2::tdT mice, 25-28 days after SE, were anesthetized with isoflurane and secured in small animal stereotaxic frame (Kopf). The scalp was injected (s.c) with bupivacaine (2mg/kg) at the site of incision. Animals also received a subcutaneous dose of carprofen (5mg/kg) and buprenorphine (Ethiqa XR at 3.25mg/kg) at start of surgery for preemptive analgesia. Two cortical screw electrodes (invivo1, Roanoke, VA) were implanted. Two additional screw electrodes on the contralateral side served as ground and reference. After 3 days of recovery, mice were connected to a tethered video-EEG monitoring system. Signals were sampled at 10 kHz, amplified (x100, 8202-SE3, Pinnacle Technologies), digitized (Powerlabs16/35, AD Instruments, Colorado Springs), and recorded using LabChart 8.0 (AD instruments). Post-SE and saline injected littermate pairs of mice underwent continuous video-EEG monitoring 24h/day for up to 14 days. Spontaneous electrographic seizures were defined as high amplitude activity, at least 3 times the standard deviation of baseline that were accompanied by simultaneous behavioral seizures greater than stage 4.

### Slice Physiology

One week after SE, experimental mice were anesthetized under isoflurane and decapitated. Mice, 1 month post SE, were perfused with ice cold sucrose ACSF under isoflurane anesthesia prior to decapitation. Whole brains were extracted, and horizontal brain slices (350 µm) were prepared using Leica VT1200S Vibratome in ice cold sucrose artificial cerebrospinal fluid (sucrose-aCSF) containing (in mM): 85 NaCl, 75 sucrose, 24 NaHCO_3_, 25 glucose, 4 MgCl_2_, 2.5 KCl, 1.25 NaH_2_PO_4_, and 0.5 CaCl. Slices were bisected and incubated at 32°C for 30 min in a holding chamber containing an equal volume of sucrose-aCSF and recording aCSF and were subsequently held at room temperature for an additional 30 min before use. The recording aCSF contained (in mM): 126 NaCl, 2.5 KCl, 2 CaCl_2_, 2 MgCl_2_, 1.25 NaH_2_PO_4_, 26 NaHCO_3_, and 10 D-glucose. All solutions were saturated with 95% O_2_ and 5% CO_2_ and maintained at a pH of 7.4 for 2-6 hours (Afrasiabi et al., 2022; Proddutur et al., 2023). Slices were transferred to a submerged recording chamber and perfused with oxygenated aCSF at 33°C. Whole-cell voltage-clamp and current-clamp recordings from GCs and presumed SGCs in the IML and outer edge of the granule cell layer were performed under IR-DIC visualization with Nikon Eclipse FN-1 (Nikon Corporation) or Nikon Eclipse NiE (Nikon Corporation) using 40x water immersion objectives. Recordings were obtained using Molecular Devices MultiClamp 700B amplifiers. Data were obtained using glass microelectrodes (3-7MΩ) pulled using Narishige PC-10 puller (Narishige Japan) or Sutter P-1000 Glass puller (Sutter Instruments) and were low pass filtered at 2kHz, digitized using Axon DigiData 1400A (Molecular Devices) or DigiData 1400A/D converter (Molecular Devices) and acquired using pClamp11 at 10kHz sampling frequency. Recordings were performed using K-gluconate internal solution containing 126 mM K-gluconate, 4 mM KCl, 10 mM HEPES, 4 mM Mg-ATP, 0.3 mM Na-GTP, and 10 mM PO-creatinine or cesium methane sulfonate (CsMeSO_4_) internal solution containing 140 mM cesium methane sulfonate, 10 mM HEPES, 5 mM NaCl, 0.2 mM EGTA, 2 mM Mg-ATP, and 0.2 mM Na-GTP (pH 7.25; 270–290 mOsm). Biocytin (0.2%) was included in the internal solution for post hoc cell identification (Yu et al., 2015; Swietek et al., 2016; Dovek et al., 2025a). Current clamp recordings to assess intrinsic active and passive parameters were obtained from SGCs and GCs, with pipettes filled with K-gluconate internal. Cells were held at −70 mV and responses to 1500 ms current steps from -200 pA to +200pA in 40 pA steps were recorded. CsMeSO_4_ based internal was used to obtain voltage clamp recordings of spontaneous and evoked EPSCs and IPSCs from a holding potential of -70 mV and 0mV, respectively. Evoked EPSCs and IPSCs were elicited by stimulation of the perforant path (PP) using a FHC stimulating electrode placed at the junction of the dorsal blade and crest, just outside the fissure (Korgaonkar et al., 2020). Iso-Flex (AMPI Israel) stimulating box was used to deliver 10 µs stimulus pulses at 0.5-2 mA intensity and 0.5 mA increase in intensity every 3 sweeps. Recordings were discontinued if series resistance increased by > 20% or if access resistance surpassed 25MΩ. Post hoc biocytin immunostaining and morphologic analysis was used to definitively identify SGCs and GCs included in this study. See “Cell Morphology” below.

### Behavioral Paradigm: The Barnes Maze

One month after SE induction, male and female pilocarpine and saline treated TRAP2::tdT mice were trained in a Barnes maze spatial learning task (Dovek et al., 2025b; Dovek et al., 2025a). The Barnes maze table (Maze Engineers, https://conductscience.com/maze/) was 92cm in diameter and consisted of 20 holes that are 5cm in diameter. There was a false floor installed with a removable escape box that can be replaced with an additional false floor piece. The maze was set up in the middle of 4 curtain walls with two bright lights and a camera for recording above the maze. Attached to each curtain was a different set of visual cues (various shapes cut from felt). The escape hole was positioned in between two of the visual cues. Animals were kept outside of the curtain in a dark room until their turn to run the trial. Animals were split into two groups: active group or “trained” that did the Barnes maze training and the control group or “untrained” that went through daily habituation but never trained in the behavior paradigm.

#### Habituation

Three days before testing, animals were handled for about a minute every day to habituate animals to the experimenter. One day (Day 0) before beginning behavior, animals were individually housed due to the hyperactive nature of this particular mouse line and in order to prevent confounding results. Animals were brought to the behavior room and left alone to habituate in their cages for at least 1 hour before beginning training. Additional habituation was performed on day 1 of training, during which animals were placed in the starter cup on the table for 1 minute. They were then guided towards the escape box (in a temporary location different from the experimental location) and led into the escape box.

#### Acquisition

For days 1-6 of training, each animal in the experimental group performed 3 trials, 180 seconds each. For those animals in the active experimental group, the experimenter performed each trial for all the animals before starting the next trial with a minimum inter-trial-interval of 15 minutes. If at the end of the 180 seconds they did not enter the escape, the experimenter guided them to the escape and then placed them back into their home cage.

#### Ensemble Induction

On day 6, the animals were brought into the room 5 hours before testing. 10 minutes before Barnes maze testing, each mouse received 4-hydroxy tamoxifen injections. Animals were left in the room for an additional 5 hours after testing to limit neuronal activity labeling not related to the behavioral paradigm.

#### Probe Trial

On day 7, the escape box was replaced to look like all the other holes. The table was rotated 180 degrees to account for potential olfactory cues. Each animal was given 90 seconds to explore the table before being returned to the home cage.

#### Reacquisition

A week following induction (6 days following probe trial) for day 12-13, animals were rerun on acquisition trials (the same as those performed on days 1-6) to reactivate the behaviorally-relevant ensemble. Animals were sacrificed for IHC 90 minutes after performing the task on day 13.

Between each trial, the entire table and escape box were wiped down with 70% ethanol and allowed to dry fully. Behavior was analyzed using Anymaze software by a blinded experimenter and using BUNS analysis software (Illouz et al., 2016; Dovek et al., 2025a).

### *4-Hydroxytamoxifen* Preparation and Administration

4-Hydroxy tamoxifen was dissolved in 100% ethanol at a concentration of 20mg/mL by sonicating solution at 37°C for 30 minutes or until it was fully dissolved. Solution was then aliquoted and stored at -20°C. On day of use, 4-OHT was redissolved by sonicating solution at 37°C for 10 minutes. A 1:4 mixture of castor oil and sunflower seed oil was added respectively to give a final concentration of 10mg/ml. The remaining ethanol in solution was evaporated by speed vacuuming in a centrifuge. Animals received 50mg/kg 4-OHT that was delivered intraperitoneally (DeNardo et al., 2019; Dovek et al., 2025b; Dovek et al., 2025a).

### Cell Morphology and Immunohistochemistry

Following physiological recordings, slices were fixed in 0.1mM phosphate buffer containing 4% paraformaldehyde at 4°C overnight. Slices were washed with PBS and then incubated in Alexa Fluor® 594 conjugated streptavidin (1:1000 Thermo Fisher, S11227) in PBS with 0.3% Triton X-100 for 2 hours at room temperature (Swietek et al., 2016; Afrasiabi et al., 2022; Dovek et al., 2025b) for cell identification.

Following Barnes maze reacquisition trials, mice and litter mate controls were euthanized with Euthasol and perfused with PBS followed by a 4% paraformaldehyde solution (Dovek et al., 2025b; Dovek et al., 2025a). The brains were left in the 4% paraformaldehyde at a temperature of 4°C for 3 hours before being transferred to PBS. The brains were sliced coronally into 50μm sections using a Leica vt100s vibratome and 5 sequential sections were chosen, each 250μm apart across the septotemporal axis and split medially for quantification. Free floating sections were blocked in 10% goat serum in PBS with 0.3% Triton X-100 for 1 hour. Sections were incubated in 4°C overnight in primary antibody for c-*Fos* (1:750, Rabbit mAb Cell Signaling Technology, cat #2250). The following day, sections were incubated in goat anti-rabbit Alexa Fluor® 488 secondary antibody (1:500 Abcam, cat #150077) for 1 hour. Slices were mounted on a glass slide using Vectashield®. Sections were visualized and imaged at 40X magnification using a Zeiss Axioscope-5 with stereo investigator imaging software for analysis. Images of TRAP2::tdT labeled cells were used to distinguish SGCs from GCs by a trained investigator blinded to experimental groups. Live images were used to verify classification as needed. The region of interest evaluated was the dentate gyrus granule cell layer (GCL) and IML. Classification was performed manually and SGCs were defined by the presence of multiple (>2) primary dendrites, wide dendritic arbor (Fig. 1I), greater soma width than height, and/or location in or close to the IML as in our earlier studies (Dovek et al., 2025b; Dovek et al., 2025a). These criteria are based on our prior studies in which unbiased cluster analysis of GC and SGC morphometric data identified the number of dendrites, soma aspect ratio and dendritic arbor width as the main factors distinguishing the cell types (Gupta et al., 2020; Afrasiabi et al., 2022). Only cells with more than 70% of their somata visible in the section were included in the counts. Any partial cells labeled with tdT, with only small portions of the soma visible, were excluded. Any cells that could not clearly be classified into one of two groups and were located in the middle or inner third of the GCL were classified as GCs.

**Figure 1:**
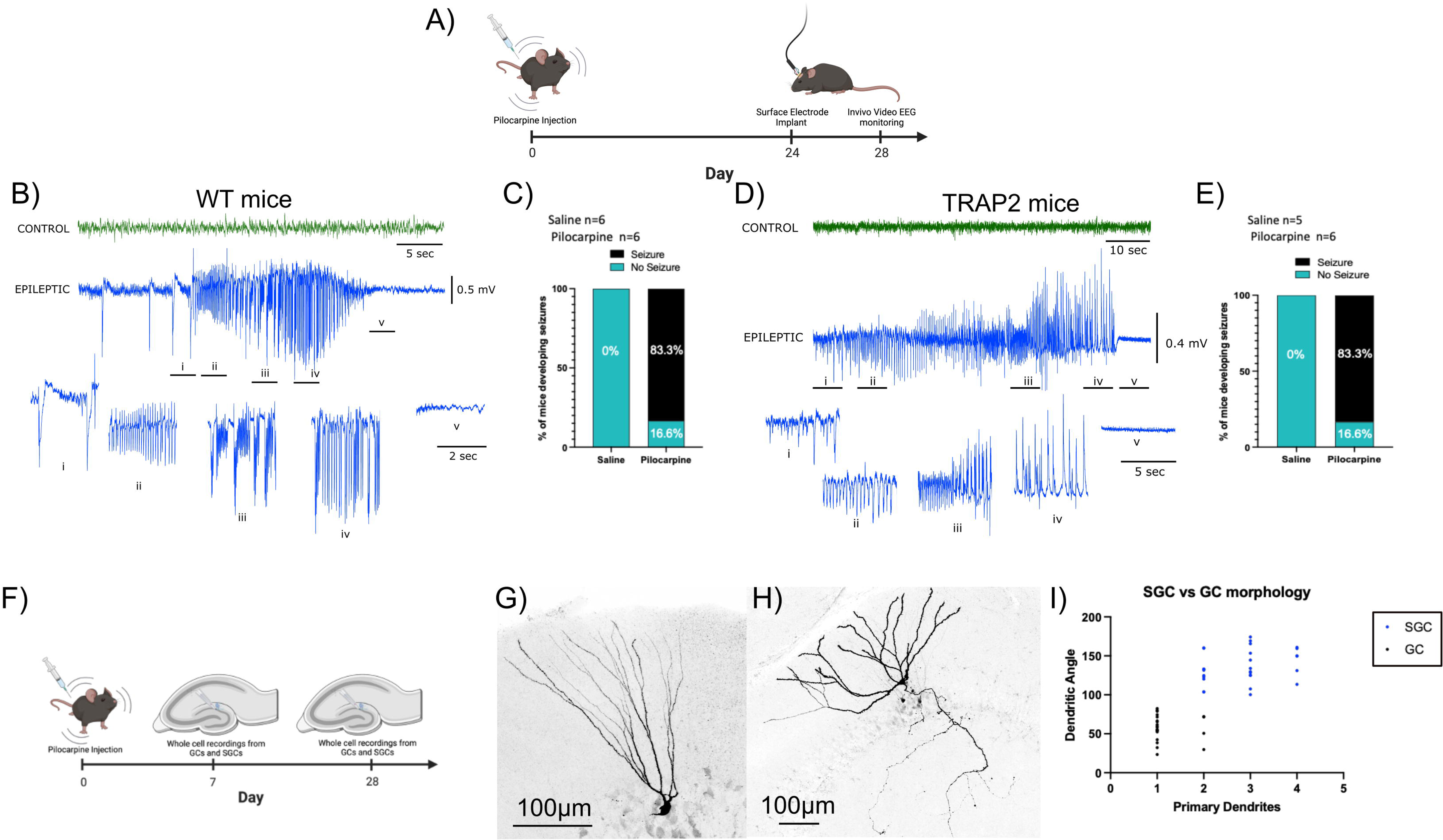
Reliable development of spontaneous recurrent seizures after pilocarpine induced status epilepticus in wild-type and TRAP2::reporter mouse lines. A) Schematic of experimental timeline for evaluation of epileptogenesis. B) Representative EEG traces from a WT mouse injected with saline (top-green) and a mouse that experienced pilocarpine SE (bottom-blue) 28 days prior to recordings. C) Histogram depicting proportion of WT animals treated with saline or pilocarpine that developed spontaneous recurrent seizures. D) Representative EEG traces from a TRAP2::reporter mouse injected with saline (top-green) and a littermate mouse that experienced pilocarpine SE (bottom-blue) 28 days prior to recordings. E) Summary plot of proportion of TRAP2::reporter animals that developed spontaneous recurrent seizures. F) Schematic of experimental timeline for slice physiology experiments. G-H) Representative images of a biocytin filled GC (G) illustrates narrow dendritic arbor while the SGC (H) has multiple primary dendrites and a wide dendritic span. I) Plot shows that a combination of number of primary dendrites and dendritic angle of biocytin filled cells can distinguish SGC from GCs.

### Quantification and Statistical Analysis

In-house custom Python scripts using eFEL package (https://github.com/BlueBrain/eFEL) were used to extract, input resistance (R_in_), resting membrane potential (RMP), action potential threshold and firing frequency (Ranjan et al., 2024). Results were validated by threshold detection analysis in Clampfit 11.

Individual spontaneous and evoked EPSCs and IPSCs were detected and analyzed using EasyElectrophysiology software (Easy Electrophysiology Ltd). A threshold search algorithm was used with events confirmed by experimenter. The first event was counted after a 180 second adjustment period. Any “noise” that spuriously met trigger specifications was rejected. Area Under the Curve (pA/ms), Rise Time (ms), and Biexponential Fit of Decay (ms) were calculated using the average trace of all events (100) per cell. For analysis of sIPSC/sEPSC frequency and amplitude 100 events per cell were used to generate cumulative probability plots and analyzed using nonparametric Kolmogorov-Smirnov (KS) Test for comparison of effect of SE on frequency and amplitude within cell type with Cohen’s D used to evaluate effect size For frequency analysis, inter-event intervals (IEIs) extracted from EasyElectrophysiology were used to compute instantaneous frequency of events as the reciprocal of the IEI. A biexponential fit was used when calculating decay. Rise time was calculated between 10 and 90% of the event amplitude. Half width is calculated as the full width at half maximum amplitude. Analyses of effect of cell type, treatments and interactions were performed on within cell averages of synaptic parameters using two-way repeated measures (RM) ANOVA, followed by pairwise multiple comparisons using Holm-Sidak method or Dunn’s method as appropriate

Sample sizes were not predetermined and conformed with those employed in the field. Significance was set to p<0.05, subject to appropriate Bonferroni Correction. Statistical analysis was performed using GraphPad Prism 10. Unpaired Mann-Whitney, two-way repeated measures (RM) ANOVA, or one-way ANOVA followed by pairwise multiple comparisons using Holm-Sidak method or Dunn’s method as appropriate. Data are presented as mean ± SEM. Additional graphics were created with BioRender.com.

## Results

### Wild-type and TRAP2::reporter mice develop spontaneous recurrent seizures one month after pilocarpine SE

In order to compare SE-induced changes in GC and SGC physiology during the early post-SE period and following development of epilepsy, we first established a time point at which experimental mice reliably developed spontaneous recurrent seizures after SE induction. Mice injected with pilocarpine or saline, were implanted with cortical screw electrodes and underwent continuous EEG-monitoring starting 4 weeks (28 days) after treatment for up to 14 days (Fig 1A-E). We observed that 83.3% (5 of 6) of WT mice that experienced SE exhibited spontaneous seizures, which were consistently accompanied by behavioral seizures greater than stage 4 on the Racine Scale (Fig 1B, C) (Racine, 1972). In contrast, none of the saline injected mice (6 of 6) displayed seizure activity. Similarly, 83.3% (5 of 6) of TRAP2::reporter mice that experienced SE and none of the saline injected controls developed spontaneous electrographic and behavioral seizures by 4-5 weeks after SE (Fig 1D, E). These data demonstrate that, in our experimental paradigm, pilocarpine-induced SE reliably leads to the development of spontaneous recurrent seizures by 1 month after SE induction. Therefore, both WT and TRAP2::reporter mice 1 month after SE, were presumed to be epileptic in subsequent experiments.

### Early and transient increase in SGC excitability after SE

To determine the effects of SE on intrinsic properties and synaptic inputs to GCs and SGCs, we conducted whole-cell-patch-clamp recordings from mice 1 week and 1 month after SE induction and in age-matched saline injected controls (Fig 1F). Recorded cells were biocytin filled and stained post-hoc to distinguish GCs and SGCs based on previously established morphological criteria (Gupta et al., 2020; Afrasiabi et al., 2022; Dovek et al., 2025b; Dovek et al., 2025a). GCs were identified based on tear-drop shaped somata in the cell layer and compact dendritic arbors with one or two primary dendrites (Fig 1G,I). SGCs were identified based on semilunar somata in or near the IML and wide dendritic arbors with two or more primary dendrites (Fig 1H,I). As with saline injected controls, GCs and SGCs in post-SE and epileptic mice were reliably distinguished based on morphology (Fig 2A, B). Interestingly, axons from both GCs and SGCs in a mouse 1-month after SE were found to have collaterals in the GCL suggestive of axonal sprouting (Fig 2B). Comparison of passive properties of the cell types in mice 1 week and 1 month after SE (Fig 2C-J) failed to reveal seizure-induced changes. However, there was an early increase in firing frequency in response to current injections in SGCs 1 week after SE (Fig 2D,H; two-way RM ANOVA, main effect of current x treatment p=0.0404). This increase in SGC excitability was not sustained and trended lower than in controls by 1 month (Fig 2F,J). Neither GCs nor SGCs showed changes in firing threshold, input resistance (R_in_) or resting membrane potential (RMP) 1 week after SE (Fig 2 K-M, Supplementary Table 1). However, GCs from mice 1-month post-SE had a more hyperpolarized action potential threshold, despite having no change in their firing frequency (Fig 2N, Supplementary Table 1, Threshold: Control GC: -31.43 ± 1.74, SE GC: -36.82± 0.79, p=0.03 by two-way RM ANOVA). Neither cell type showed changes in R_in_ or RMP 1 month after SE (Fig 2O, P). Collectively, these data identify an early increase in SGC excitability that recovers by 1 month and a delayed lowering of action potential threshold in GCs which suggests that the cell types develop distinct post-SE changes in intrinsic physiology that contribute to network instability after SE.

**Figure 2:**
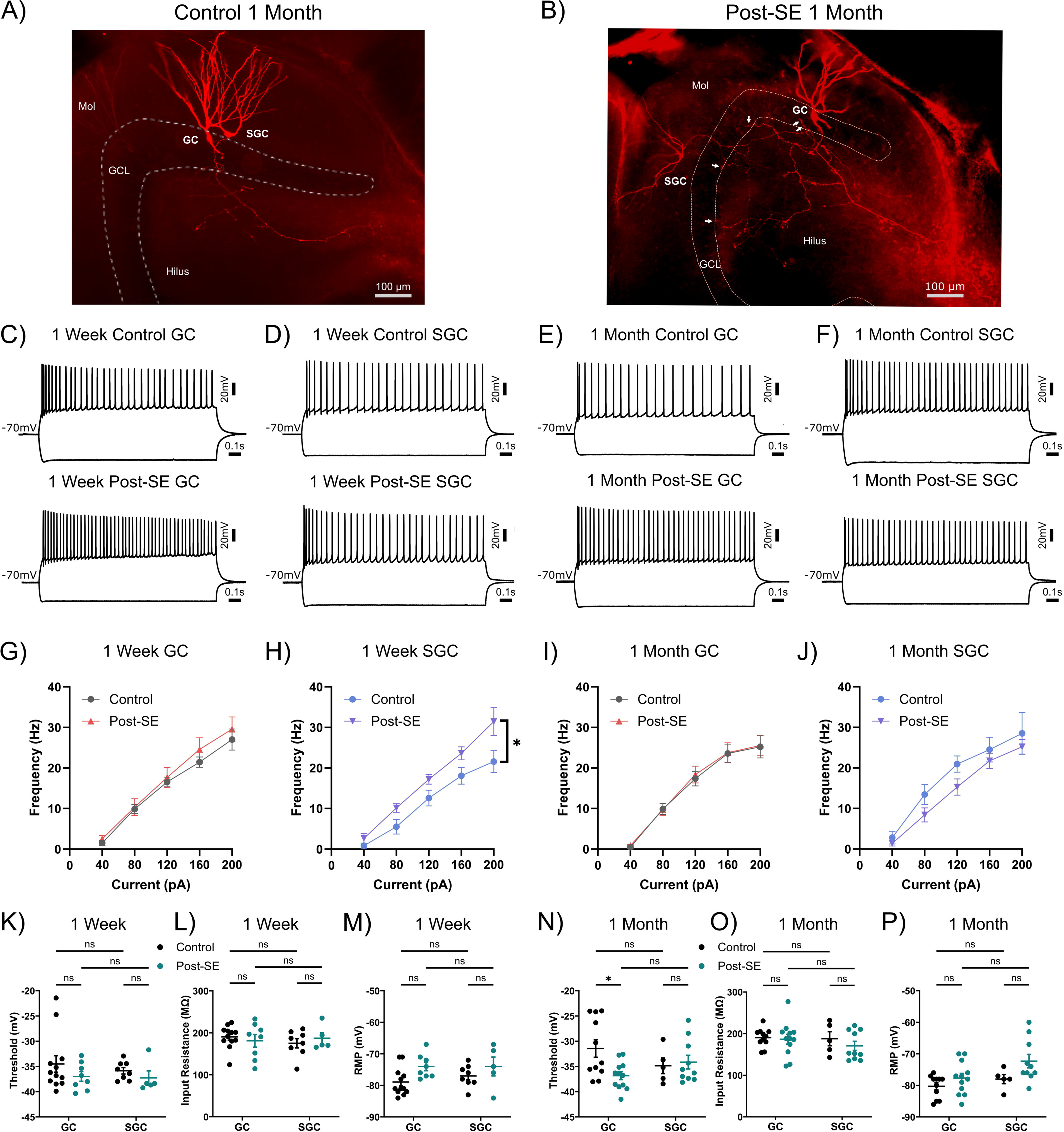
Early and selective increase in SGC excitability after SE. A-B) Representative biocytin fills of a GC and SGC from a control animal (A) and a mouse 1-month after SE (B) obtained following dual patch clamp recordings. White arrows denote recurrent axon collaterals. C-D) Representative cell membrane voltage traces in response to +160 pA and - 200 pA current injections in 1 week control GC (top, C), and post-SE GC, (bottom, C), control SGC (top, D) and post-SE SGC (bottom, D). E-F) Representative cell membrane voltage traces in response to +160 pA and -200 pA current injections in 1 month control GC (top, E), GC 1-month post-SE, (bottom, E), control SGC (top, F) and SGC 1-month post-SE (bottom, F). G-H) Summary plot of firing frequency in response to increasing current injections in GCs (G) and SGCs (H) 1 week following saline (control) or pilocarpine (post-SE) injections. * indicates p<0.05 Two-way ANOVA, main effect of current x treatment p=0.0404). I-J) Summary plot of firing frequency in response to increasing current injections in GCs (I) and SGCs (J) 1 month following saline or pilocarpine injections. K-M) Summary plots of action potential threshold (K), input resistance (L) and resting membrane potential (M) between cell types and treatments 1 week after SE. N-P) Summary plots of threshold (N), input resistance (O), and resting membrane potential (P) between cell types and treatments 1 month after pilocarpine/saline injections. * indicates p<0.05. Data were obtained from n=cells/ mice: 1-week Control: 12 GCs / 4 mice and 8 SGCs / 4 mice; 1-week post-SE: 8 GCs / 4 mice and 5 SGCs / 3 mice; 1-month Control: 11 GCs/ 4 mice and 5 SGCs / 4 mice; 1-month post-SE: 12 GCs / 3 mice and 10 SGCs / 4 mice.

### Early and sustained increases in spontaneous excitatory inputs to SGCs and GCs after SE

Although GCs have been extensively studied in SE and found to exhibit abnormal sprouting of axon collaterals increase in excitatory synaptic inputs, whether SGCs show similar changes is unknown. We find that both GCs and SGCs show an increase in sEPSC frequency with a moderate effect size 1 week after SE (Fig 3A-D; p<0.05 by K-S test Cohen’s d= 0.37 for GC and 0.42 for SGC). Consistently, analysis of sEPSC frequency averaged within cell revealed a significant effect of cell type and treatment with both cell types demonstrating an increase in sEPSC frequency (Supplemental Figure 1, Supplementary Table 2, two-way RM ANOVA, main effect of cell type p<0.0001, treatment p<0.0001 and interaction p=0.015). While sEPSC amplitude in GCs showed a small decrease with low effect size after SE (Fig 3E,F; p<0.05 by K-S test, Cohen’s d=0.19 for GC), sEPSC amplitude in SGCs remained unchanged. Accordingly, cell averages of sEPSC amplitude failed to show effects of either cell type or treatment (Supplemental Figure 1, Supplementary Table 2, by two-way RM ANOVA). Although there were cell type and treatment effects on decay time, which were consistent with faster decay in SGCs compared to GCs and after SE compared to controls, pairwise comparisons within cell type failed to reach statistical significance (Supplementary Table 2; two-way RM ANOVA, main effect of cell type p=0.03, main effect of treatment p=0.04). No significant differences were observed in rise time and charge transfer.

**Figure 3:**
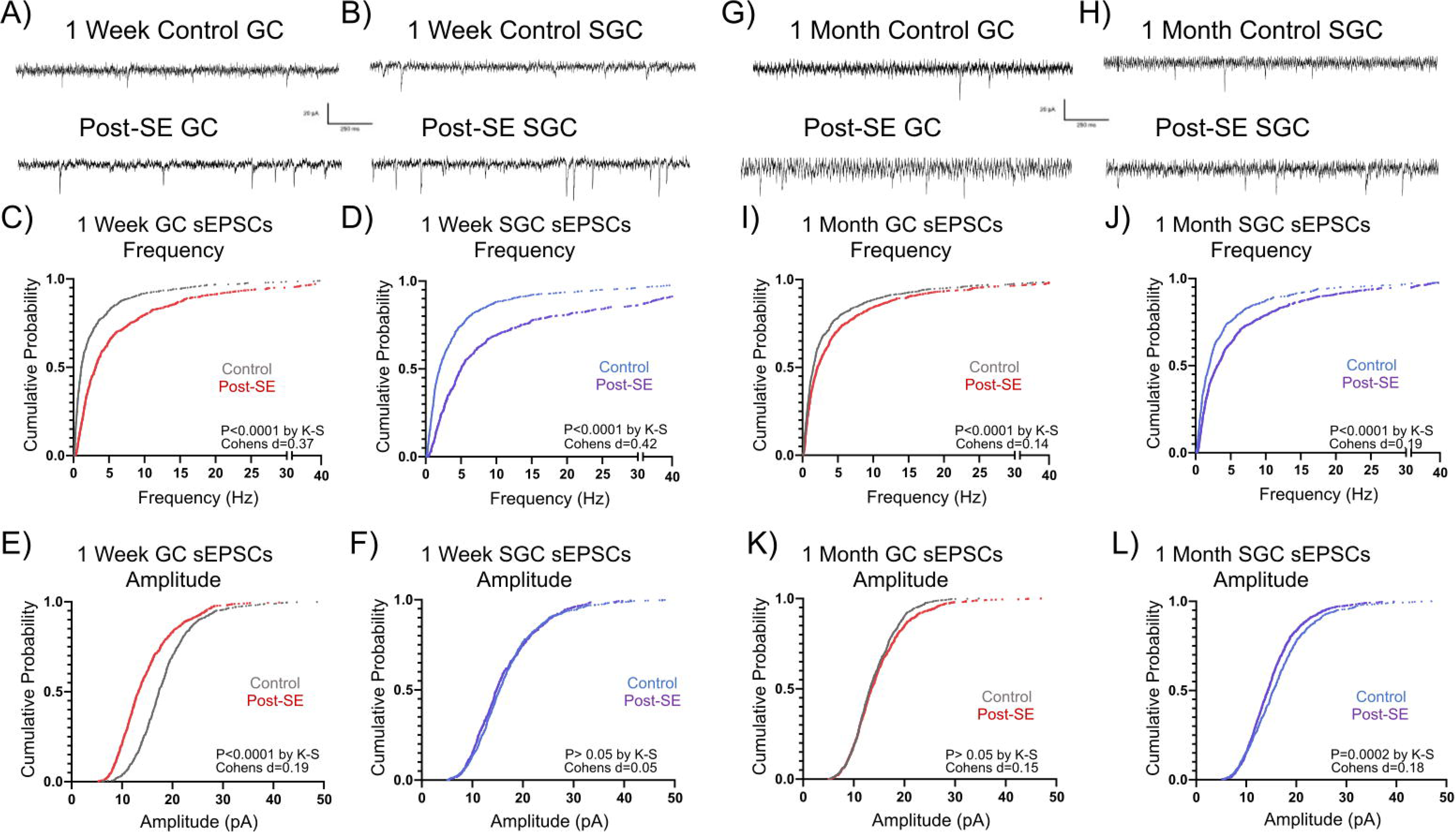
Early and sustained increase in frequency of spontaneous excitatory inputs in both GCs and SGCs. A-B) Representative current traces from GCs and SGCs show sEPSCs recorded 1 week after SE induction in control GC (A-top), post-SE GC (A-bottom), control SGCs (B-top), and SGCs post-SE (B-bottom). C-D) Cumulative probability plots compare sEPSC frequency between post-SE and control GCs (C) and SGCs (D). (E-F) Cumulative probability plots compare sEPSC amplitude between post-SE and control GCs (E) and SGCs (F). Data were obtained from n=cells/ mice: 1-week Control: 8 GCs / 4 mice and 9 SGCs / 6 mice; 1-week post-SE: 8 GCs / 5 mice and 6 SGCs / 4 mice. G-H) Representative current traces from GCs and SGCs show sEPSCs 1 month following SE control GC (G-top), GCs 1-month post-SE (G-bottom), control SGCs (H-top) and SGCs 1-month post-SE (H-bottom). I-J) Cumulative probability plots comparing sEPSC frequency between 1-month post-SE and control GCs (I) and SGCs (J). (K-L) Cumulative probability plots comparing amplitudes of sEPSCs between 1-month post-SE and control GCs (K) and SGCs (L). Data were obtained from n=cells/ mice: 1-month Control: 10 GCs / 5 mice and 7 SGCs / 5 mice; 1 month post-SE: 10 GCs / 5 mice and 10 SGCs / 6 mice. *indicates p<0.05 TW-ANOVA with Šídák’s multiple comparisons post hoc tests.

Similarly, sEPSC frequency in both GCs and SGC remained higher 1 month after SE with a modest effect (Fig 3G-J; p<0.05 by K-S test, Cohen’s d=0.14 in GC and 0.19 in SGC). Analysis of the within cell averages of sEPSC frequency revealed a significant effect of both cell type and treatment (Supplemental Figure 1, Supplementary Table 3, two-way RM ANOVA, main effect of cell type and p=0.0001, treatment p<0.0001). Specifically, the sEPSC frequency in control SGCs was higher than in control GCs and both GCs and SGCs showed an increase in sEPSC frequency 1-month after SE (Supplemental Figure 1, Supplementary Table 3). In contrast, while sEPSC amplitude in GCs was not different from controls by 1 month, SGCs develop a decrease in sEPSC amplitude with small effect size 1 month after SE (Fig 3K,L; by K-S test Cohen’s d=0.18 in SGC). Cell averaged sEPSC amplitude in controls trended higher in SGCs than in GCs (Supplemental Figure 1, Supplementary Table 3, main effect of cell type p=0.055 for control GC vs SGC) with no effect of treatment on either cell type. By 1 month post pilocarpine/saline injection, there were no effects of treatment on sEPSC decay, rise time or charge transfer in either cell type (Supplementary Table 3). However, there was a significant effect of cell type on charge transfer with sEPSCs in SGCs exhibiting higher charge transfer than in GCs (Supplementary Table 3: two-way RM ANOVA, main effect of cell type p=0.032). These findings demonstrate that both GCs and SGCs undergo an early and sustained increase in sEPSC frequency and trends towards temporally dynamic cell-type specific changes in sEPSC amplitude after SE.

### Early and transient increase in SGC sIPSC frequency after SE

Examination of inhibitory synaptic inputs identified a main effect of pilocarpine-SE on sIPSC frequency and revealed a modest early increase in sIPSC frequency in SGCs but not in GCs (Fig 4 A-D, Supplementary Table 4; K-S test, Cohen’s d=0.03 in GC and 0.30 in SGC). Correspondingly, sIPSC frequency averaged within cell revealed a significant effect of treatment with a selective increase in sIPSC frequency in SGCs after SE (Supplemental Figure 2, Supplementary Table 4, two-way RM ANOVA, main effect of treatment p<0.05). There was an early post-SE increase in sIPSC amplitude with small effect in GCs and SGCs (Fig 4E-F, by K-S test Cohen’s d = 0.29 in GC and 0.05 in SGC). While this trend remained in cell averaged data, it did not reach statistical significance (Supplemental Figure 2, Supplementary Table 4, two-way RM ANOVA, main effect of treatment p=0.06). There was no effect of cell-type or treatment on sIPSC charge transfer or biexponential decay time in either GCs or SGCs (Supplementary Table 4). However, while GCs showed no change in sIPSC rise time after SE, there was a significant decrease in sIPSC rise time in SGCs after SE (Supplementary Table 4; two-way RM ANOVA, effect of treatment p=0.0017). These findings raise the possibility of differential inhibitory innervation of GCs and SGCs after SE.

**Figure 4:**
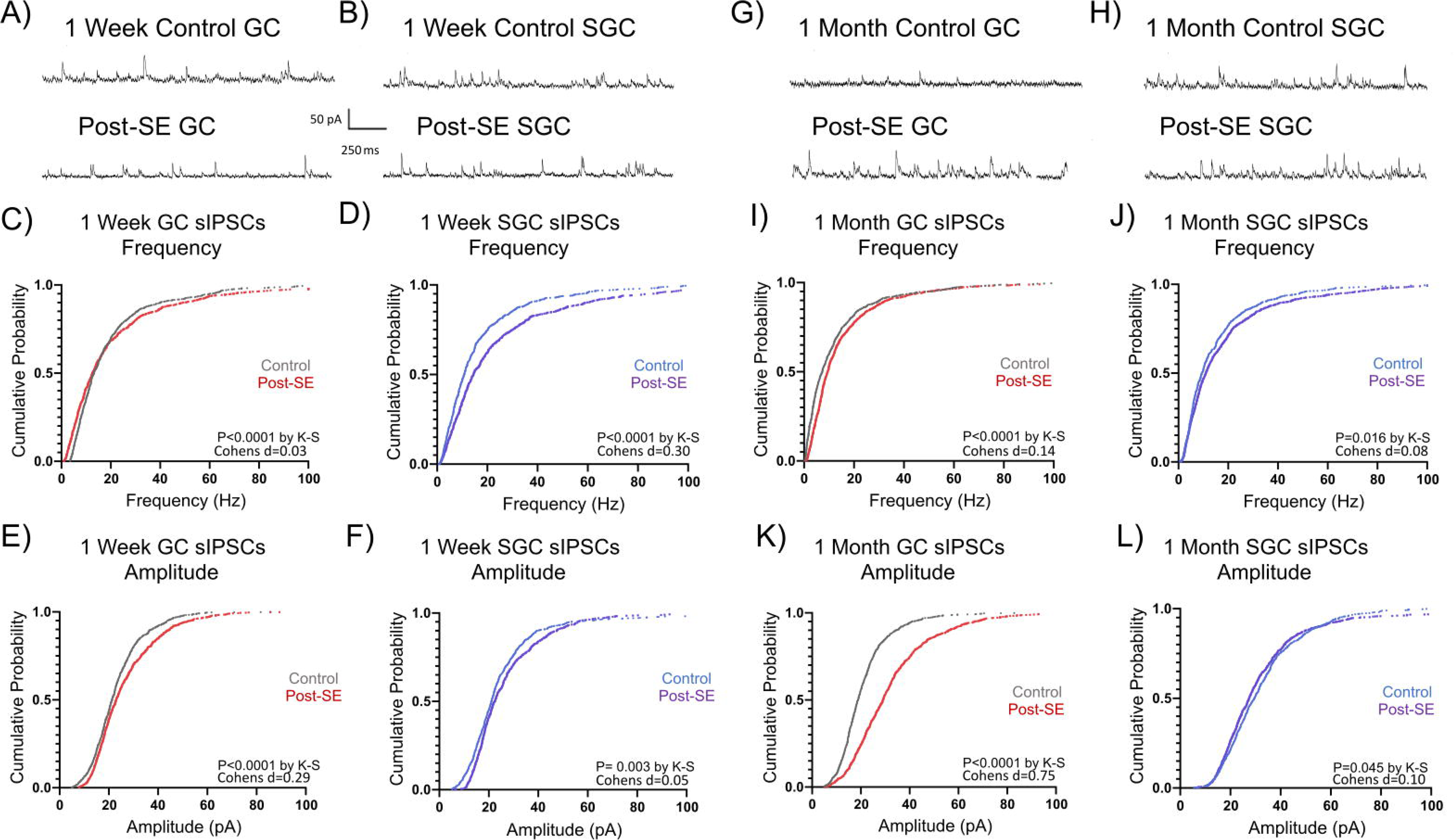
Early transient increase in sIPSC frequency in SGCs after SE. A-B) Representative current traces from GCs and SGCs show sIPSCs 1 week after SE in control GC (A-top), post-SE GC (A-bottom), control SGCs (B-top), and post-SE SGCs (B-bottom). C-D) Cumulative probability plots compare sIPSC frequency between post-SE and control GCs (C) and SGCs (D). (E-F) Cumulative probability plots compare sIPSC amplitude between post-SE and control GCs (E) and SGCs (F). Data were obtained from n=cells/ mice: 1-week Control: 10 GCs / 5 mice and 9 SGCs / 5 mice; 1-week post-SE: 9 GCs / 5 mice and 7 SGCs / 4 mice. G-H) Representative current traces from GCs and SGCs show sIPSCs 1 month following SE induction in control GC (G-top), GCs 1-month post-SE (G-bottom), control SGCs (H-top) and SGCs 1-month post-SE (H-bottom). I-J) Cumulative probability plots compare sIPSC frequency between 1-month post-SE and control GCs (I) and SGCs(J). (K-L) Cumulative probability plots of sIPSC amplitude in epileptic and control GCs (K) and SGCs (L) Data were obtained from n=cells/ mice: 1-month Control: 9 GCs / 5 mice and 9 SGCs / 5 mice; 1 month post-SE: 11 GCs / 6 mice and 10 SGCs / 6 mice. *indicates p<0.05 TW-ANOVA with Šídák’s multiple comparisons post hoc tests.

In mice 1 month after SE, sIPSC frequency in both GCs and SGCs was different from corresponding sham controls but the effect size was low (Fig 4G-J, K-S test, Cohen’s d= 0.14 in GC and 0.08 in SGC). Analysis of cell averaged data revealed no post-SE change in sIPSC frequency in either cell type at one month (Supplemental Figure 2, Supplementary Table 5). However, GCs showed a robust post SE increase in sIPSC amplitude (Fig 4K-L, by K-S test,, Cohen’s d=0.75) and in cell averaged data (Supplemental Figure 2, Supplementary Table 5, by two-way RM ANOVA, effect of treatment p=0.02), which was also associated with an increase in sIPSC charge transfer in GCs (Supplementary Table 5). SGCs showed no change in amplitude or charge transfer in mice 1-month post-SE (Supplementary Table 5). SGCs continued to have faster sIPSCs rise times in mice 1-month after SE, with main effects of treatment and interaction between cell type and treatment on sIPSC rise times (Supplementary Table 5; two-way RM ANOVA, effect of treatment p=0.04 and effect of interaction p=0.016). This differential regulation suggests that SGCs are subject to distinct inhibitory control mechanisms during epileptogenesis, potentially reflecting their unique role in network dynamics and memory encoding.

### SGCs show distinct early and delayed changes in perforant path driven inputs in experimental epilepsy

The perforant path provides the primary excitatory input to the dentate gyrus, and its activity is tightly regulated by feedforward and feedback inhibitory circuits critical for controlling network excitability and shaping information flow. To determine if SE alters afferent driven input and inhibition in GCs and SGCs, we compared perforant path evoked EPSC and IPSC parameters between the cell types. At 1 week after SE, no significant differences in evoked EPSC (eEPSC) amplitude were observed between cell types or as a result of treatment (Fig 5A-D, Supplementary Table 6). However, by 1 month post-SE, a significant decrease in eEPSCs was observed specifically in SGCs (Fig 5 E-H, Supplementary Table 6). On the other hand, eIPSC amplitude showed significant effect of SE and cell-type 1 week after SE with a significant decrease in eIPSC amplitude in SGCs (Fig 5I-L, Supplementary Table 7). However, these differences were no longer apparent 1 month after SE, suggesting circuit level compensation over time (Fig 5M-P, Supplementary Table 7). Despite these changes, the excitation inhibition ratio (E/I ratio) measured as the ratio of the peak amplitude of the eEPSC to the eIPSC within the same cell was not different in either cell type in mice 1-week or 1-month after or epileptic mice (Figure 5Q-R). Thus, despite the multifaceted circuit level changes after SE, the E/I balance in DG projection neurons appears to be maintained.

**Figure 5:**
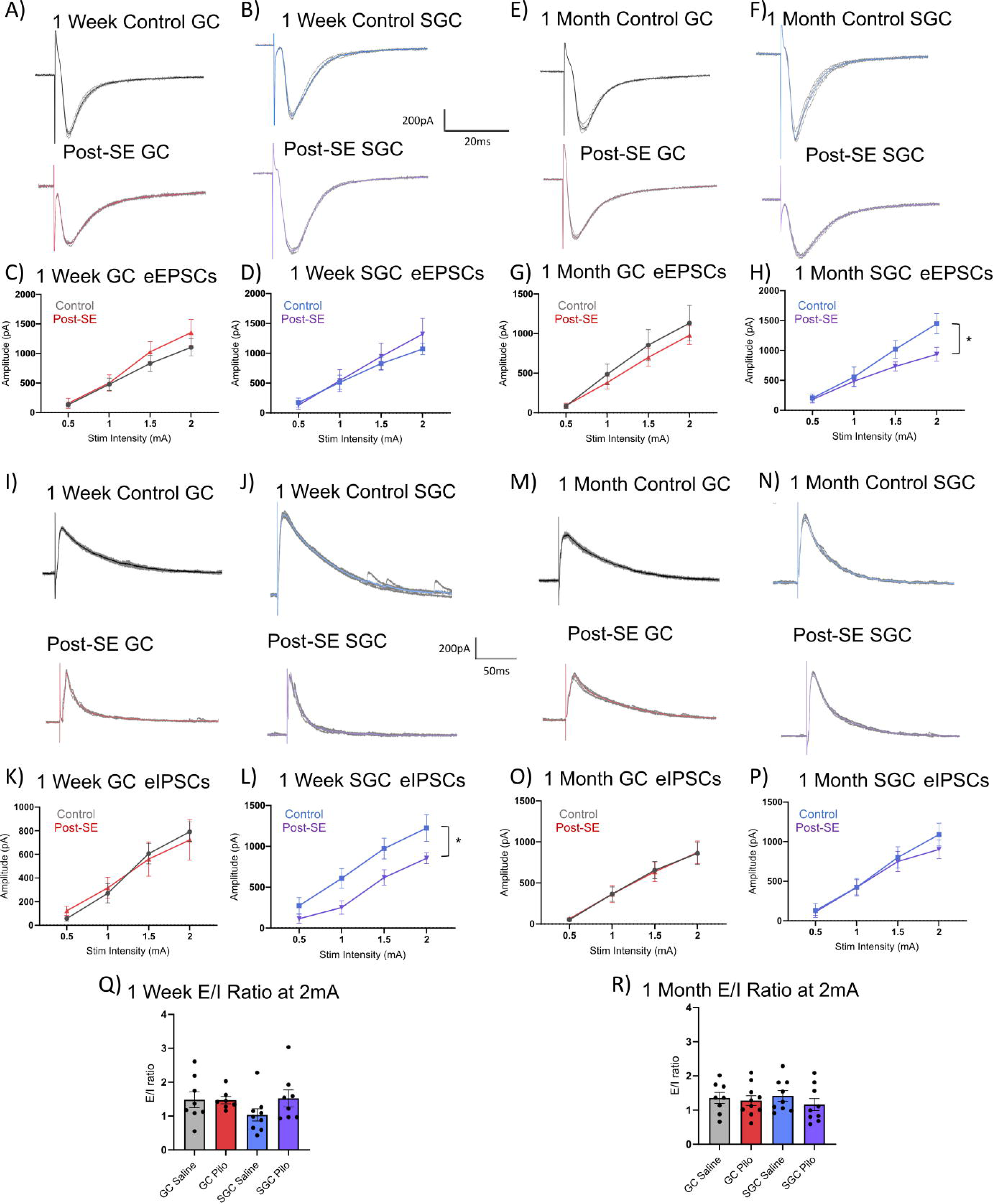
Selective and time-dependent alterations in perforant path evoked synaptic events in SGCs. A-B) Representative perforant path evoked EPSCs in a control GC (A, top), post-SE GC (A, bottom), control SGC (B, top), and post-SE SGC (B, bottom) 1 week after treatment. C-D) Summary plot of evoked EPSC peak amplitude in response to PP stimuli at increasing intensities in GCs (C) and SGCs (D) 1 week after SE. Data were obtained from n=cells/ mice: 1-week Control: 8 GCs / 5 mice and 9 SGCs / 6 mice; 1-week post-SE: 7 GCs / 5 mice and 8 SGCs / 5 mice. E-F) Representative perforant path evoked EPSCs in a control GC (E, top), GC 1-month post-SE (E, bottom) control SGC (F, top) and SGC 1-month post-SE (F, bottom). G-H) Summary plot of evoked EPSCs at increasing stimuli intensities in GCs (G) and SGCs (H) 1 month following treatment. Data were obtained from n=cells/ mice: 1-month Control: 8 GCs / 5 mice and 9 SGCs / 5 mice; 1 month post-SE: 10 GCs / 6 mice and 9 SGCs / 6 mice. I-J) Representative perforant path evoked IPSCs in a control GC (I, top), post-SE GC (I, bottom), control SGC (J, top) and post-SE SGC (J, bottom) 1 week after treatment. K-L) Summary plot of evoked IPSCs at increasing stimuli intensities in GCs (K) and SGCs (L) 1 week after saline or pilocarpine injections. Data were obtained from n=cells/ mice: 1-week Control: 8 GCs / 5 mice and 9 SGCs / 6 mice; 1-week post-SE: 7 GCs / 5 mice and 8 SGCs / 5 mice. M-N) Representative perforant path evoked IPSCs in a control GC (M, top), post-SE GC (M, bottom) control SGC (N, top) and post-SE SGC (N, bottom) 1 month after treatment O-P) Summary plot of evoked IPSCs at increasing stimuli intensities in GCs (O) and SGCs (P) 1 month following treatment. Data were obtained from n=cells/ mice: 1-month Control: 8 GCs / 5 mice and 9 SGCs / 5 mice; 1 month post-SE: 10 GCs / 6 mice and 9 SGCs / 6 mice. Q-R) Summary plot of E/I ratio at 2 mA stimulation 1 week (Q) and 1 month (R) after saline or pilocarpine injections.

### Spatial navigation is impaired in mice after SE

To test how epilepsy impacts spatial learning, male and female TRAP2::tdT mice underwent pilocarpine induced SE and were trained in the Barnes maze task 1 month later when they were presumed epileptic. Littermate saline and pilocarpine treated task naïve behavioral controls (untrained mice) were brought to the testing room with mice undergoing training but were never trained on the maze. The untrained mice served as control to assess basal labeling of neurons independent of behavioral task in TRAP2::tdT mice following our tamoxifen induction protocol. During the initial acquisition phase of the Barnes maze task, both experimental (trained) groups showed significant improvements in task performance as evident from the decrease in primary latency to find the escape during the first few days. The treatment group effect on primary latency trended towards but failed to reach statistical significance (p=0.0640), but the main effect was seen in primary latency across acquisition days with animals in both groups requiring less time to first encounter the escape (two-way RM ANOVA p<0.0001). These data demonstrate that post-SE mice learn the task. However, post-SE mice made more errors before finding the escape zone (two-way RM ANOVA, main effect of treatment p=0.018), suggesting that the post-SE mice may adopt different strategies to perform the task or may have impaired spatial awareness. We evaluated spatial awareness through analysis of search strategy using BUNS analysis software (Illouz et al., 2016; Dovek et al., 2025b) (Fig 6D-F), an automated tool that categorizes animal’s search strategy as random, serial search (sequential search), or spatial (showing direct or corrected pathways) based on tracking analysis from video recordings. While control animals increasingly adopt more spatial search strategies over acquisition trials, post-SE mice continue to use random or serial search approaches to locate the escape box (Fig 6D-E). To quantify these search strategies, each strategy was associated with a cognitive score from “0” for random to “1” for direct spatial strategies (Illouz et al., 2016; Dovek et al., 2025b). The average cognitive scores of post-SE mice diverged significantly from control mice by acquisition day 4, with control mice having much higher cognitive score indicating use of spatial strategy (Fig 6F; two-way RM ANOVA, main effect of treatment p<0.0001). Task performance and use of spatial cues to orient themselves was also examined during a probe trial. The proportion of the total 90 second test time during which the animal (both trained and untrained groups) stayed within the quadrant where the escape used to be located was quantified as an assessment of spatial recall (Fig. 6 B-F probe trial). To limit potential confounds, the table was turned by180 degrees prior to the probe trial to isolate involvement of visual cues guiding spatial search strategy. Trained post-SE mice exhibited increase in primary latency and errors (Fig 6 B-C), reduced use of spatial strategy (Fig 6 D-E) and reduced cognitive score during the probe trials than trained control mice. Time spent by trained control and post-SE mice in escape quadrant was compared with corresponding untrained littermate groups. Trained control mice spent significantly more time in the escape zone compared to both untrained groups and the trained post-SE mice (Fig 6G; two-way RM ANOVA, main effect of treatment p<0.0001, main effect of training condition p=0.0004, and a main effect of interaction p=0.002), suggesting that only saline injected trained mice were able to use spatial cues to search for the escape zone. These data identify that although post-SE mice appear to learn the task, their use of spatial navigation strategies and recall based on spatial cues is significantly compromised.

**Figure 6:**
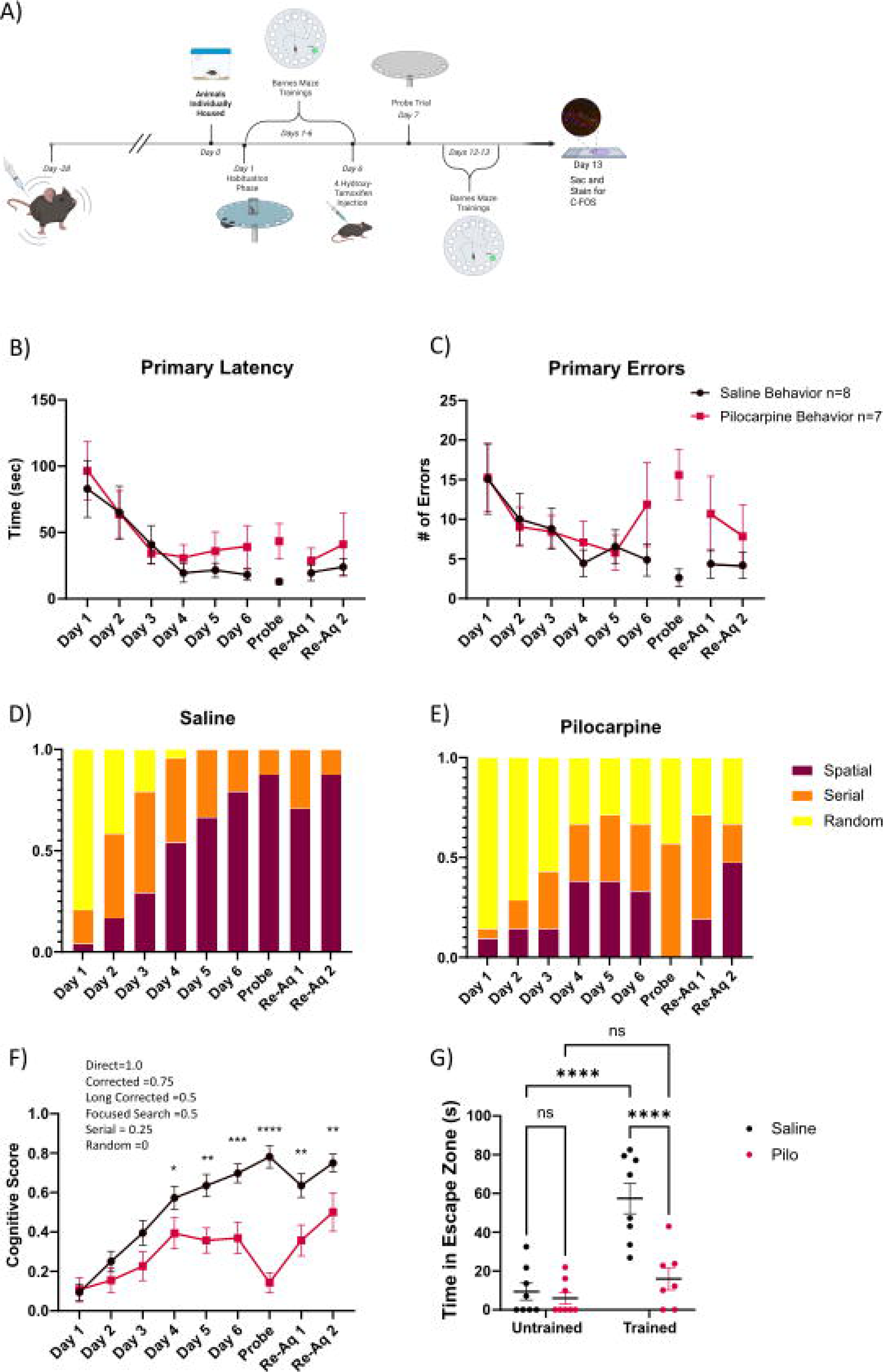
Mice are impaired in the use of spatial search strategy in a Barnes Maze spatial navigation task after SE. A) Schematic of experimental timeline for pilocarpine or saline injections. B-C) Summary plots of primary latency to first inspection of the escape hole (B) and number of errors until first exit inspection (C) in trained saline and post-SE mice. D-E) Plot of search strategies used in trained control (D) and post-SE (E) over the course of experimental days. F) Quantification of cognitive score determined by search strategy used in trained saline and post-SE mice. G) Summary plot of overall time spent in escape quadrant during the probe trial in untrained and trained saline and post-SE mice. *indicates p<0.05, ** indicates p<0.01, *** indicates p<0.001, **** indicates p<0.0001 by TW-ANOVA with Šídák’s multiple comparisons post hoc tests. N= 8 untrained mice each, 8 trained control and 7 trained post-SE mice.

### Altered ensemble size and SGC recruitment after SE

To test how epilepsy impacts task-related recruitment of SGCs and GCs to activity dependent neuronal ensembles, we induced tdT labeling of neurons in TRAP2::tdT mice on day 6 of the Barnes maze acquisition, when control mice were found to reliably use spatial strategy to locate the escape box, and in task-naïve littermate controls (Fig 6A, Fig 7A-D). Initial analysis of untrained, task-naïve mice identified a significant increase in tdT labeled neurons in post-SE mice suggesting a basal increase in excitability (Fig 7B,D,E untrained control 5.34 ± 0.45; untrained post-SE 8.98 ± 2.1, p=0.02 by Mann Whitney). In contrast, trained post-SE mice had fewer labeled tdT neurons than trained saline-controls (Fig 7A,C,E; trained control 1.60 ± 1.20, trained post-SE: 9.2± 1.66 p=0.0006 Mann-Whitney). To test the reliability of reactivation of the tdT-labeled neuron during recall, mice were sacrificed 90 minutes after a second re-acquisition trial (Re-aq 2 indicated in Fig. 6) and immunostained for c-*Fos*. Predictably, untrained saline-injected control and post-SE mice had fewer neurons which colocalized tdT and immunostained c-*Fos* (Fig 7F). Interestingly, untrained control animals showed more co-labeling compared to untrained post-SE mice. (Fig 7F; control untrained:0.03± 0.01, epileptic untrained: 0.0066± 0.004, p=0.025). Consistent with ensemble reactivation during recall, trained saline-treated and post-SE mice had significantly more neurons co-labeled for tdT and c-*Fos* than in corresponding task naïve mice. However, trained post-SE mice had significantly fewer neurons which co-labeled for tdT and c-*Fos* than trained saline-controls (Fig 7F; Multiple Mann Whitney tests, trained control: 0.08 ± 0.015; untrained control: 0.03 ± 0.01 p= 0.0004; trained control vs trained post-SE: 0.03± 0.009 p=0.0012). Further *post hoc* scrutiny of data revealed two outliers with extremely high tdT labeling density in task-naive and trained post-SE groups. To evaluate the possibility that the outlier data may have resulted from occurrence of electrographic seizures during the Cre induction window, we compared c-*Fos* immunostaining in mice euthanized 90 minutes after induction of an acute pilocarpine SE to that of saline-controls (Fig 7G,H). DG c-*Fo*s expression levels in sections from post-SE mice which were outliers for c-*Fo*s labeling density were qualitatively similar to c*-Fos* expression in sections from mice after acute SE (Fig. 7H-I) consistent with occurrence of seizures within the induction window. Based on the strategy adopted in prior studies (Godale et al., 2022), we reasoned that mice with tdT expression well over 2 standard deviations above the rest of their cohorts (Fig 7 I-J) had seizures close to or during the Cre induction window and excluded data from these two mice in subsequent analysis. With removal of the outliers, untrained post-SE mice had significantly *fewer* tdT labeled neurons than in untrained saline-controls (Fig 7K, untrained control: 5.34± 0.45, untrained post-SE: 2.76 ± 0.43 p<0.0001 Mann-Whitney test). Similarly, trained post-SE mice had *fewer* tdT labeled neurons than trained saline controls (Fig 7K, trained control: 11.60 ± 1.2, trained post-SE: 5.54 ± 0.72 p<0.0001 Mann-Whitney test). These data identify an overall decrease in basal and behaviorally relevant neuronal activity in post-SE mice in the absence of an acute seizure episode. However, both trained control and post-SE mice had significantly more tdT labeled neurons than their respective untrained counterparts demonstrating recruitment of neurons to behaviorally relevant ensembles after training (Fig 7K, trained vs untrained control p<0.0001, trained vs untrained post-SE p=0.0023 by multiple Mann-Whitney tests). As expected based on task dependent neuronal activation, trained saline-injected mice had significantly more neurons colocalizing tdT and c-*Fos* following re-acquisition training (Fig 7L trained control:0.08± 0.05, untrained control: 0.03 ± 0.01 p=0.0004; multiple Mann-Whitney tests), while the small increase in co-labeling in trained post-SE mice compared to untrained post-SE mice failed to reach statistical significance (trained vs untrained post-SE: 0.0073± 0.005 p=0.058 Mann-Whitney test). Notably, co-labeling of tdT with c-*Fos* was significantly reduced in trained post-SE mice compared to trained saline-controls (Fig 7L; trained control vs trained post-SE: 0.037 ± 0.02 p=0.0002). The increased co-localization of tdT and c-*Fos* in trained saline-controls compared to untrained counterparts is consistent with tdT labeling after training representing task-related neurons which are, consequently, more reliably reactivated following reacquisition. In contrast, post-SE mice have a significant reduction in basal activity levels in untrained mice and a relatively blunted increase in activity levels during training.

**Figure 7:**
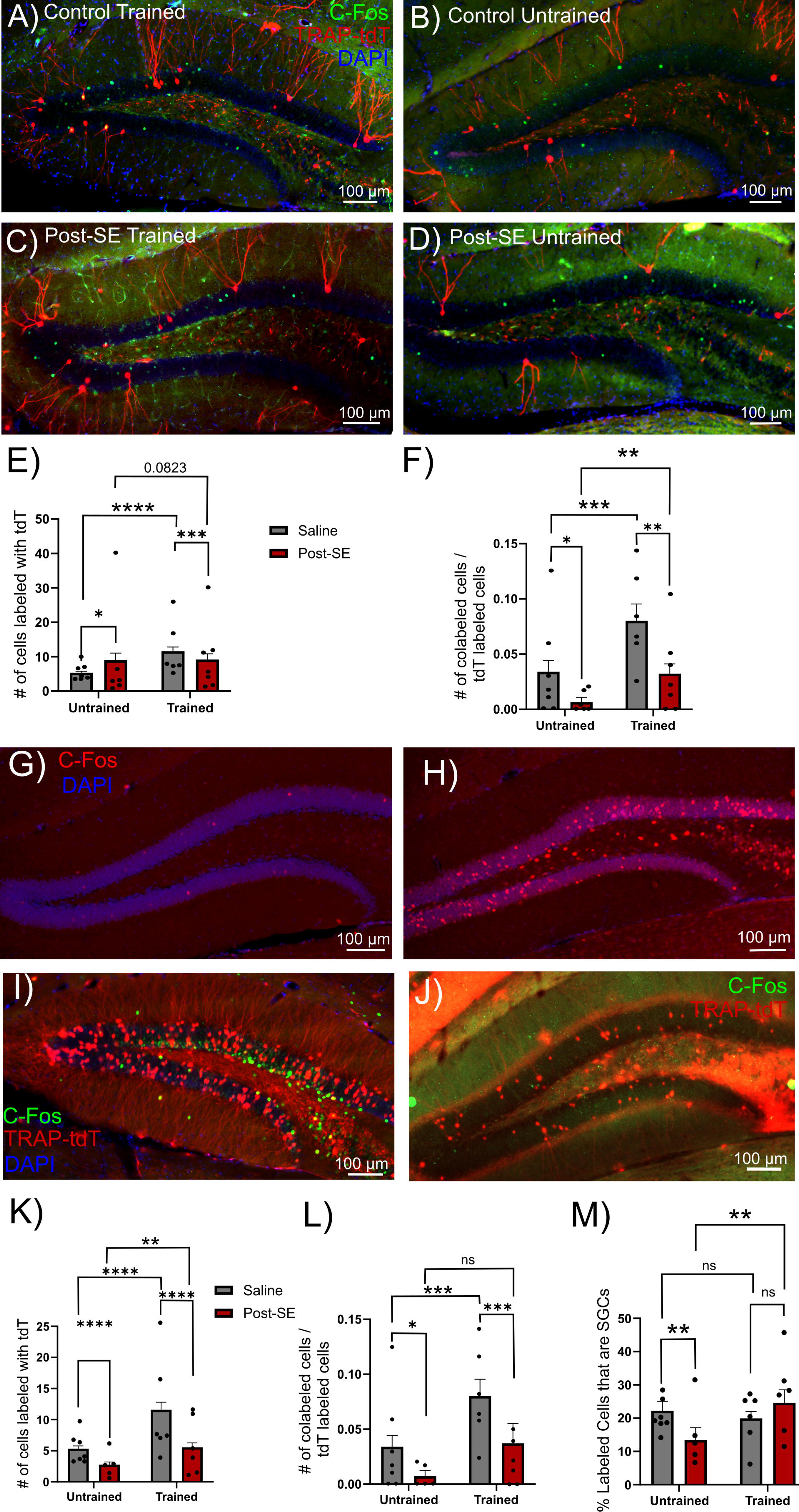
Altered proportional recruitment of SGCs to basal and task-related DG ensembles in post-SE mice. A-D) Representative epifluorescence image of sections from TRAP2::tdT mice one week after cre induction on day 6 of acquisition illustrates tdT labeling (red) and subsequent labeling for *c-Fos* immunostaining (green) in saline trained (A), saline littermate untrained control mice (B), pilocarpine injected trained and SE littermate untrained post-SE mice (D). E-F) Summary of number of tdT labeled cells/section (E) and the proportion of tdT positive cells co-labeled for *c-Fos* (green) (F) in 9-10 sections each from 7 untrained control, 6 untrained post-SE, 6 trained control and 7 trained post-SE mice. G-H) Representative epifluorescence images of *c-Fos* immunostaining (red) for immediate early gene expression 90 minutes after saline injection (G) and acute pilocarpine induced SE (H). Data points represent within animal averages of slices used in analysis. I-J) Representative epifluorescence image of sections from the two post-SE mice that likely had a seizure during the “trapping” window based on tdT labeling levels higher than 2 SD over average. K-M) Summary of number of tdT labeled cells/ section (K) and proportion of tdT positive cells co-labeled for *c-Fos* (green) (L) with the mice in panels I and J excluded for analysis due to suspected seizure withing Cre induction window. M) Quantification of percentage of tdT labeled cells in K that had morphology consistent with SGCs *indicates p<0.05, ** indicates p<0.01, *** indicates p<0.001, **** indicates p<0.0001 by Šídák’s multiple comparisons post hoc tests, in 9-10 sections each from 7 untrained control, 5 untrained post-SE, 6 trained control and 6 trained post-SE mice.

Finally, to determine if SGC recruitment to active neuronal ensembles is altered after SE, we quantified tdT labeled SGCs and GCs based on morphological criteria (Fig 1) (Gupta et al., 2020; Dovek et al., 2025b; Dovek et al., 2025a). Consistent with our previous findings, the proportion of tdT labeled SGC was 20-25% in both untrained and trained saline-control mice (Fig 7M, trained control: 19.92 ± 2.06, untrained control: 22.21 ± 2.8 p=0.95, Mann-Whitney test). Crucially, the proportion of tdT labeled SGCs in untrained post-SE mice was significantly lower than in untrained controls suggesting reduced basal SGC activity after SE. However, proportional recruitment of SGCs among tdT labeled neurons in trained post-SE mice was similar to that of trained saline injected mice (Fig 7M; trained control: 19.92 ± 2.06; trained post-SE: 24.62 ± 3.9, p=0.77, Mann-Whitney test). Notably, there was an increase in the proportional recruitment of SGCs following training in post-SE mice (Fig7M, untrained post-SE: 13.42 ± 3.73, trained post-SE: 24.62 ± 3.9, p=0.0025, Mann-Whitney test) which was not observed in controls (Fig7M, untrained control: 22.21 ± 2.8, trained control: 19.92 ± 2.06, p>0.05, Mann-Whitney test). Together, the tdT labeling data identify a reduction in basal SGC recruitment to neuronal ensembles in after SE indicating reduced SGC basal activity and an increase in task dependent SGC recruitment which could contribute to altered DG spatial memory processing in epilepsy.

## Discussion

The dentate gyrus has a central role in regulating hippocampal excitability and information flow. Following brain insults and during epileptogenesis, the hippocampal circuitry undergoes extensive changes. In this study, we focused on SGCs, an understudied DG projection neuron subtype with sustained firing whose alterations during epileptogenesis are unknown. Our data (summarized in Figure 8) identify that SGCs develop an early increase in intrinsic excitability and a decrease in afferent evoked inhibitory current amplitude. These changes have the potential to enhance DG excitability and throughput. Although both SGCs and GCs received more excitatory inputs after SE, SGCs also received more frequent spontaneous inhibitory synaptic inputs, suggesting cell-type specific differences in circuit connectivity and plasticity. SGC show a unique early increase in excitability after SE which returns to levels observed in controls by 1 month. However, the action potential threshold in GCs is hyperpolarized 1-month after SE, which could support increased dentate excitability. DG neuronal recruitment to basal and task driven ensembles was diminished in mice 1-month after SE suggesting a paradoxical reduction in activity and/or c-*Fos* activation in epilepsy. Moreover, *c-Fos* colocalization in tdT labeled neurons following task reacquisition was decreased after SE. This post-SE decrease in colocalization of *c-Fos* expression, in response to task re-acquisition, with neuronal ensembles recruited during task learning was associated with impaired use of spatial search strategies. Interestingly, proportional SGC recruitment to active neuronal ensembles under task-naïve basal conditions was reduced in after SE but was not different from controls following task exposure. These data suggest that task-dependent recruitment of SGCs is maintained in epilepsy. Collectively, these findings identify that SGCs undergo a distinct set of intrinsic and synaptic alterations during epileptogenesis that could shape both disease progression and spatial navigation.

**Figure 8:**
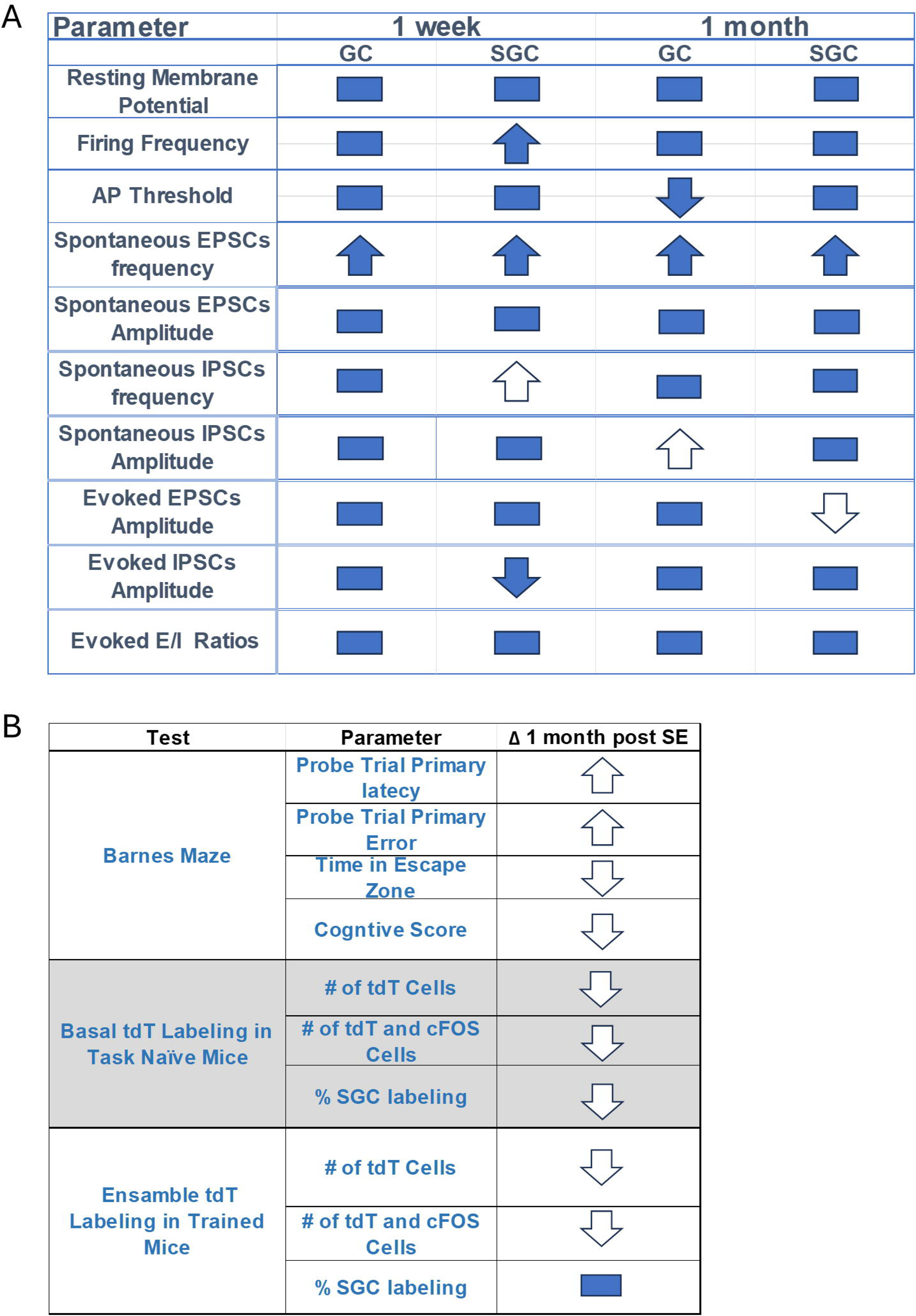
Summary of cell-type specific physiological changes as well as alterations in behavioral and activity-dependent ensemble labeling after SE. Table A. Summary of the significant changes in various parameters in GCs and SGCs compared to corresponding controls. Changes that are expected to increase excitability are represented by filled arrows. Changes that decrease excitability are represented as open arrows. Arrow direction indicates direction of change from corresponding control. Table B. Summary of the significant changes in behavioral test and activity-dependent ensemble labeling between groups.

The changes in SGC intrinsic excitability and synaptic inputs occur early after SE and are distinct from those observed in GCs. Uniquely, SGCs show greater firing in response to depolarization early after SE. Because SGCs have axon terminals in CA3, it is interesting to speculate that increase in SGC firing could promote CA3 excitability and contribute to epileptogenesis. Conversely, since SGC firing is linked to enhanced feedback inhibition in the DG (Larimer and Strowbridge, 2010; Afrasiabi et al., 2022), it is equally likely that the increase in SGC excitability may develop as a compensatory mechanism to recruit dentate inhibition. In mice 1-month post-SE, GCs exhibited a more hyperpolarized action potential threshold without change in firing frequency suggesting potential homeostatic compensation to maintain GC activity levels. Together, both early and delayed cell-type specific post-SE changes in GC and SGC intrinsic physiology have the potential to enhance network excitability.

The increase in sEPSC frequency observed in SGCs at 1 week and 1 month after SE is consistent with that reported in GCs (Althaus et al., 2019). The frequency of sEPSC in SGC in control mice was greater than in GCs, as reported previously (Dovek et al., 2025a), and increased further after SE. The increase in excitatory input frequency following SE could result from development of recurrent excitatory mossy fiber collaterals (Buckmaster et al., 2002; Shibley and Smith, 2002). Indeed, it is possible that the sprouted collaterals provide powerful drive to SGCs by targeting spines observed on SGCs somata in the IML (Dovek et al., 2025a). Unlike sEPSC frequency which showed similar changes in both cell types after SE, sEPSC amplitude trended to be reduced in GCs, but not SGCs. The enhanced spontaneous excitatory drive to SGCs after SE could augment dentate up states and network excitability (Larimer and Strowbridge, 2010). Curiously, the amplitude of perforant path driven EPSCs is decreased in SGCs from mice 1-month after SE suggesting that these complementary changes may serve to differentially recruit SGCs under different activity states. While the increase in SGC excitability may reflect an attempt to enhance feedback inhibition, it is possible the net effect of increasing SGC excitability in the epileptic DG is pro excitatory due to the loss of their hilar interneuron targets which support feedback inhibition (Buckmaster and Dudek, 1997; Kapur, 2003; Larimer and Strowbridge, 2010; Afrasiabi et al., 2022). Molecular access to selectively manipulate SGCs is currently lacking and needed to directly evaluate SGC contribution to dentate excitability in healthy and epileptic circuits.

Previous studies have identified that SGCs and GCs differ in developmental and injury-induced changes in tonic and synaptic inhibition (Gupta et al., 2012; Gupta et al., 2022). Here we report that, while GCs in mice 1 week and 1-month post SE show no change in sIPSC frequency, SGCs develop an early transient increase in sIPSC frequency after SE. Moreover, although consistent with previously reported increase in postsynaptic GABA_A_ receptor densities in GCs (Brooks-Kayal et al., 1998; Kobayashi and Buckmaster, 2003), sIPSC amplitude in GCs is increased in mice 1-month post-SE, it remains unchanged in SGCs. Furthermore, there is an early post-SE decrease in evoked IPSC amplitude selectively in SGCs which recovers to control levels 1 month after SE. These cell-type specific changes in sIPSC frequency and amplitude during epileptogenesis support dynamic and divergent circuit level compensation for interneuron loss (Houser and Esclapez, 1996; Zhang et al., 2009; Butler et al., 2022). Interestingly, the sIPSC rise times in SGCs, but not GCs, is considerably faster after SE than in controls, suggesting that SGCs receive more perisomatic inputs compared to GCs (Soltesz et al., 1995). This differential perisomatic innervation is supported by our previous finding of greater parvalbumin interneuron mediated feedback inhibitory inputs to SGCs during synaptic barrages (Afrasiabi et al., 2022). These findings of cell-type specific changes in inhibition during epileptogenesis extend current knowledge on differences in inhibitory circuit connectivity and plasticity between GCs and SGCs.

Having identified multiple cell type specific changes in GCs and SGCs following SE and since SGCs show preferential labeling as part of cellular memory engrams (Erwin et al., 2020; Dovek et al., 2025b), we used TRAP::tdT mice to evaluate SGC recruitment to behaviorally-relevant neuronal ensembles in mice1 month after SE. Despite needing a higher pilocarpine dose for SE induction, TRAP2::reporter mice reliably developed spontaneous seizures 1 month after SE. Although the post-SE TRAP2::tdT mice showed no deficits in task acquisition indicating that the mice learned the task, the probe trials revealed a decrease in time spent in the region of the escape, suggesting deficits in use of spatial cues for navigation. A potential reason for the discrepancy between “learning” during task acquisition and “recall” in probe trial may be due to averaging of 3 trials on acquisition days allowing for learning within the day while the probe trial was a single attempt. Indeed, mice 1-month after SE consistently showed improved performance within the same day during acquisition trials (not shown). Nevertheless, we cannot exclude the possibility that post-SE mice exhibited interictal spiking during behavioral testing or developed altered arousal, locomotion or anxiety which impacted task performance during the probe trial. Unexpectedly, when excluding the two mice with excessive labeling suggestive of seizure activity from analysis, post-SE mice had fewer tdT labeled neurons in both task-naïve and trained conditions. It is possible that disruption of circadian rhythm associate with the 10 hours of dark housing prior to tamoxifen induction predisposed some mice to seizures (Quigg, 2000; Liu et al., 2022). Exclusion of mice with c-*Fos* labeling consistent with seizures was justified due to the extensive overlap between memory and seizure circuits which could disrupt memory performance (Naik et al., 2021). The decrease in basal and task-related c-*Fos* dependent neuronal labeling is likely a consequence of the multiple circuit level changes in the DG that, together, limit basal excitability of projection neurons. Alternatively, the decrease in tdT labeling in post-SE mice could occur from enhanced chronic Δ*FosB* labeling which is known to suppress c-*Fos* expression and neuronal excitability (Bing et al., 1997; Morris et al., 2000; You et al., 2017; Lamothe-Molina et al., 2022; Stephens et al., 2024). Additionally, c-*Fos* colocalization upon task re-acquisition in neurons labeled during task acquisition was reduced after SE. While these findings suggests that neuronal ensembles are less reliably reactivated during the same task after SE, it is equally possible that post-SE mice show reduced *c-Fos* induction (Stephens et al., 2024). Alternatively, post-SE mice could adopt different cognitive strategies and, consequently, use non-overlapping neuronal ensembles to perform the task during learning and re-acquisition. Collectively, the decrease in neuronal activation and reduced reactivation of task-related neuronal ensembles, could undermine recall of spatial information in epilepsy.

Although SGCs represent approximately 5% of the GC population (Save et al., 2019), over 20% of tdT labeled neurons in both trained and task naive control mice were morphologically classified as SGCs (Dovek et al., 2025b). Unexpectedly, despite their increases RMP and enhanced basal synaptic excitation, task naïve post-SE mice show a reduction in the proportion of tdT labeled SGCs compared to saline injected controls. It is interesting to speculate that reduced activation of SGCs in task naïve epileptic mice may serve to limit DG excitability and up-states. Notably, although the strength of evoked EPSCs was reduced in SGCs in mice 1-month after SE, proportional labeling of SGC in task-related DG ensembles was not different between control and post-SE mice. These results identify state-dependent difference in SGC recruitment to neuronal ensembles in post-SE mice and suggest that SGC activity may scale based on the behavioral conditions to regulate global DG excitability while retaining memory ensembles.

In summary, our results reveal that post SE changes in SGCs differ from that in GCs and identify divergent evolution of intrinsic and circuit plasticity over time during epileptogenesis. The early post-SE transient increase in SGC intrinsic excitability could promote epileptogenesis. Our findings suggest that physiological changes in SGCs in epilepsy may serve to limit SGC basal activity while retaining task-related activation and identify SGCs as a unique and critical node for plasticity during epileptogenesis.

## Acknowledgements

We thank Drs. Edward Zagha and Sachiko Haga-Yamanaka for thoughtful discussions and Dr. Mahboubeh Ahmadi for timely assistance with updating figures. We thank Dr. David Carter at the UCR Microscopy Core and Mr. Erick Contreras for help with imaging.

## Declaration of generative AI and AI-assisted technologies in the writing process

No generative AI and AI-assisted technologies were used in writing this manuscript.

