## Supplemental Table for "Altered Physiology and Ensemble Recruitment of Dentate Gyrus Semilunar Granule Cells in a Mouse Model of Epilepsy"

**Laura Dovek[1a], Andrew Huang[1] and Vijayalakshmi Santhakumar[1]**
[1]Department of Molecular, Cell and Systems Biology, University of California Riverside, Riverside, California 92521.

### Supplemental Table 1

| Figure | Parameter (Unit) | Control GC (GC-S) n=12 / 4 mice | Post-SE GC (GC-P) n=8 / 4 mice | Control SGC (SGC-S) n=8 / 4 mice | Post-SE SGC (SGC-P) n=5 / 3 mice | F & P value | Multiple Comparisons |
|---|---|---|---|---|---|---|---|
| 2 | 1 Week Resting Membrane Potential (mV) | -78.92 ± 1.27 | -74.00 ± 1.34 | -77.00 ± 1.23 | -74.00 ± 1.27 | Interaction: F (1, 29) = 0.347 P=0.5601 | GC-S vs GC-P p=0.1252 |
| | | | | | | Cell Type: F (1, 29) = 0.3474 P=0.5601 | SGC-S vs SGC-P p=0.8175 |
| | | | | | | **Treatment: F (1, 29) = 5.928 P=0.0213** | GC-S vs SGC-S p=0.9269 |
| | | | | | | by Two way ANOVA | GC-P vs SGC-P p>0.9999 |
| 2 | 1 Week Action Potential Threshold (mV) | -34.50 ± 1.640 | -36.98 ± 0.96 | -35.85 ± 0.64 | -37.26 ± 1.40 | Interaction: F (1, 29) = 0.1330 P=0.7180 | GC-S vs GC-P p=0.7102 |
| | | | | | | Cell Type: F (1, 29) = 0.3089 P=0.5826 | SGC-S vs SGC-P p=0.9909 |
| | | | | | | Treatment: F (1, 29) = 1.760 P=0.1950 | GC-S vs SGC-S p=0.9772 |
| | | | | | | by Two way ANOVA | GC-P vs SGC-P p>0.9999 |
| 2 | 1 Week Input Resistance (MΩ) | 190.3 ± 7.73 | 181.0 ± 15.07 | 175.3 ± 10.96 | 187.2 ± 12.95 | Interaction: F (1, 29) = 0.8045 P=0.3771 | GC-S vs GC-P p=0.9899 |
| | | | | | | Cell Type: F (1, 29) = 0.1363 P=0.7147 | SGC-S vs SGC-P p=0.9883 |
| | | | | | | Treatment: F (1, 29) = 0.0125 P=0.9118 | GC-S vs SGC-S p=0.9007 |
| | | | | | | by Two way ANOVA | GC-P vs SGC-P p=0.9997 |
| **Figure** | **Parameter (Unit)** | **Control GC (GC-S) n=11 / 4 mice** | **Post-SE GC (GC-P) n=12 / 3 mice** | **Control SGC (SGC-S) n=5 / 4 mice** | **Post-SE SGC (SGC-P) n=10 / 4 mice** | **F & P value** | **Multiple Comparisons** |
| 2 | 1 Month Resting Membrane Potential (mV) | -80.27 ± 1.09 | -77.67 ± 1.5 | -78.00 ± 1.45 | -72.30 ± 2.13 | Interaction: F (1, 34) = 0.7862 P=0.3815 | GC-S vs GC-P p=0.7856 |
| | | | | | | **Cell Type: F (1, 34) = 4.793 P=0.0355** | SGC-S vs SGC-P p=0.2544 |
| | | | | | | **Treatment: F (1, 34) = 5.666 P=0.0230** | GC-S vs SGC-S p=0.9584 |
| | | | | | | by Two way ANOVA | GC-P vs SGC-P p=0.1062 |
| 2 | 1 Month Action Potential Threshold (mV) | -31.43 ± 1.74 | -36.82 ± 0.79 | -34.86 ± 1.55 | -34.14 ± 1.35 | **Interaction F (1, 34) = 4.261 P=0.0467** | **GC-S vs GC-P p=0.0295** |
| | | | | | | Cell Type: F (1, 34) = 0.0636 P=0.8024 | SGC-S vs SGC-P p=0.9998 |
| | | | | | | Treatment: F (1, 34) = 2.486 P=0.1241 | GC-S vs SGC-S p=0.6185 |
| | | | | | | by Two way ANOVA | GC-P vs SGC-P p=0.6340 |
| 2 | 1 Month Input Resistance (MΩ) | 190.4 ± 6.74 | 186.6 ± 11.85 | 187.8 ± 16.93 | 170.6 ± 11.00 | Interaction: F (1, 34) = 0.3236 P=0.5732 | GC-S vs GC-P p>0.9999 |
| | | | | | | Cell Type: F (1, 34) = 0.6124 P=0.4393 | SGC-S vs SGC-P p=0.9352 |
| | | | | | | Treatment: F (1, 34) = 0.7880 P=0.3809 | GC-S vs SGC-S p>0.9999 |
| | | | | | | by Two way ANOVA | GC-P vs SGC-P p=0.8661 |

### Supplemental Table 2

| Figure | Parameter (Unit) | Control GC (GC-S) n=8/ 4mice | Post-SE GC (GC-P) n=8 / 5 mice | Control SGC (SGC-S) n=9 / 6 mice | Post-SE SGC (SGC-P) n=6/ 4mice | F & P value | Multiple Comparisons |
|---|---|---|---|---|---|---|---|
| 3 | 1 Week sEPSC Frequency (Hz) | 0.9±0.27 | 2.36±0.38 | 1.44 ± 0.26 | 3.66±0.7 | **Interaction: F (1, 27) = 6.697 P=0.015** | **GC-S vs GC-P p<0.0001** |
| | | | | | | **Cell Type: F (1, 27) = 38.55 P<0.0001** | **SGC-S vs SGC-P p<0.0001** |
| | | | | | | **Treatment: F (1, 27) = 153.4 P<0.0001** | **GC-S vs SGC-S p=0.0116** |
| | | | | | | **by Two way ANOVA** | **GC-P vs SGC-P p<0.0001** |
| | 1 Week sEPSC Amplitude (pA) | 16.26 ± 0.27 | 12.24 ± 0.60 | 13.27±1.12 | 13.28 ± 0.89 | Interaction: F (1, 27) = 1.223 P=0.2786 | GC-S vs GC-P P=0.1216 |
| | | | | | | Cell Type: F (1, 27) = 0.009687 P=0.9223 | SGC-S vs SGC-P P=0.9937 |
| | | | | | | Treatment: F (1, 27) = 1.197 P=0.2835 | GC-S vs SGC-S p=0.4579 |
| | | | | | | by Two way ANOVA | GC-P vs SGC-P p=0.4251 |
| | 1 Week sEPSCs Biexp fit Decay (ms) | 5.42 ± 0.29 | 4.73 ± 0.43 | 4.67 ± 0.65 | 3.21 ± 0.39 | Interaction:F (1, 27) = 0.5817 P=0.4523 | GC-S vs GC-P p=0.7812 |
| | | | | | | **Cell Type: F (1, 27) = 5.224 P=0.0303** | SGC-S vs SGC-P p=0.1970 |
| | | | | | | **Treatment: F (1, 27) = 4.703 P=0.0391** | GC-S vs SGC-S p=0.7080 |
| | | | | | | **by Two way ANOVA** | GC-P vs SGC-P p=0.1861 |
| | 1 Week sEPSC Area Under the Curve (pA ms) | 59.25 ± 15.21 | 49.18 ± 6.21 | 59.26 ±8.66 | 49.82 ± 6.19 | Interaction: F (1, 27) = 0.0009385 P=0.9758 | GC-S vs GC-P p=0.9286 |
| | | | | | | Cell Type: F (1, 27) = 0.0009865 P=0.9752 | SGC-S vs SGC-P p=0.9522 |
| | | | | | | Treatment: F (1, 27) = 0.8991 P=0.3514 | GC-S vs SGC-S p>0.9999 |
| | | | | | | by Two way ANOVA | GC-P vs SGC-P p>0.9999 |
| | 1 Week sEPSC Rise Time (ms) | 1.23 ± 0.07 | 1.25 ± 0.04 | 1.18 ± 0.04 | 1.20 ± 0.07 | Interaction: F (1, 27) = 0.007936 P=0.9297 | GC-S vs GC-P p=0.9997 |
| | | | | | | Cell Type: F (1, 27) = 1.011 P=0.3236 | SGC-S vs SGC-P p=0.9977 |
| | | | | | | Treatment: F (1, 27) = 0.1012 P=0.7528 | GC-S vs SGC-S p=0.8872 |
| | | | | | | by Two way ANOVA | GC-P vs SGC-P p=0.9563 |

#### Supplemental Table 3

| Figure | Parameter (Unit) | Control GC (GC-S) n=10 / 5 mice | Post-SE GC (GC-P) n=10 / 5 mice | Control SGC (SGC-S) n=7 / 5 mice | Post-SE SGC (SGC-P) n=10 / 6 mice | F & P value | Multiple Comparisons |
|---|---|---|---|---|---|---|---|
| 3 | 1 Month sEPSC Frequency (Hz) | 1.11± 0.24 | 1.62± 0.37 | 1.56 ± 0.17 | 2.15 ± 0.45 | Interaction: F (1, 33) = 0.1131, P=0.7388 | **GC-S vs GC-P p=0.0016** |
| | | | | | | **Cell Type: F (1, 33) = 19.35 P=0.0001** | **SGC-S vs SGC-P p=0.0011** |
| | | | | | | **Treatment: F (1, 33) = 24.64, P<0.0001** | **GC-S vs SGC-S p=0.0098** |
| | | | | | | by Two way ANOVA | **GC-P vs SGC-P p=0.0013** |
| | 1 Month sEPSC Amplitude (pA) | 10.18 ± 1.23 | 10.88 ±2.89 | 12.33 ± 1.01 | 11.60 ± 0.55 | Interaction:F (1, 33) = 0.9582 P=0.3348 | GC-S vs GC-P p=0.4808 |
| | | | | | | **Cell Type: F (1, 33) = 3.860 P=0.0579** | SGC-S vs SGC-P p=0.5045 |
| | | | | | | Treatment: F (1, 33) = 0.0004217 P=0.9837 | **GC-S vs SGC-S p=0.0552** |
| | | | | | | by Two way ANOVA | GC-P vs SGC-P p=0.4685 |
| | 1 Month sEPSC Biexp fit Decay (ms) | 3.91 ± 0.28 | 4.66 ±0.32 | 4.03 ±0.27 | 3.79 ±0.28 | Interaction: F (1, 33) = 3.039 P=0.0906 | GC-S vs GC-P p=0.2138 |
| | | | | | | Cell Type: F (1, 33) = 1.809 P=0.1878 | SGC-S vs SGC-P p=0.9657 |
| | | | | | | Treatment: F (1, 33) = 0.7966 P=0.3786 | GC-S vs SGC-S p=0.9980 |
| | | | | | | by Two way ANOVA | GC-P vs SGC-P p=0.1076 |
| | 1 Month sEPSC Area Under the Curve (pA ms) | 28.11 ±4.80 | 44.45 ±5.93 | 56.73± 12.34 | 46.71 ±5.33 | Interaction: F (1, 33) = 3.651 P=0.0648 | GC-S vs GC-P p=0.3057 |
| | | | | | | **Cell Type: F (1, 33) = 5.009 P=0.0321** | SGC-S vs SGC-P p=0.8032 |
| | | | | | | Treatment: F (1, 33) = 0.2103 P=0.6496 | **GC-S vs SGC-S p=0.0334** |
| | | | | | | by Two way ANOVA | GC-P vs SGC-P p=0.9987 |
| | 1 Month sEPSC Rise Time (ms) | 1.16 ±0.07 | 1.09 ±0.04 | 1.06 ±0.05 | 1.09 ±0.05 | Interaction: F (1, 33) = 0.9736 P=0.3310 | GC-S vs GC-P p=0.7794 |
| | | | | | | Cell Type: F (1, 33) = 0.9736 P=0.3310 | SGC-S vs SGC-P p=0.9905 |
| | | | | | | Treatment: F (1, 33) = 0.1486 P=0.7023 | GC-S vs SGC-S p=0.5735 |
| | | | | | | by Two way ANOVA | GC-P vs SGC-P p>0.9999 |

### Supplemental Table 4

| Figure | Parameter (Unit) | Control GC (GC-S) n=10/ 5 mice | Post-SE GC (GC-P) n=9/ 5 mice | Control SGC (SGC-S) n=9/ 5 mice | Post-SE SGC (SGC-P) n=7/ 4 mice | F & P value | Multiple Comparisons |
|---|---|---|---|---|---|---|---|
| 4 | 1 Week sIPSCs Frequency (Hz) | 9.209± 1.36 | 10.03 ±2.06 | 8.99± 1.99 | 11.78± 2.45 | Interaction: F (1, 31) = 2.192 P=0.1488 | GC-S vs GC-P p=0.3663 |
| | | | | | | Cell Type:F (1, 31) = 1.329 P=0.2579 | **SGC-S vs SGC-P p=0.0079** |
| | | | | | | **Treatment: F (1, 31) = 7.377 1 P=0.0107** | GC-S vs SGC-S p=0.8093 |
| | | | | | | by Two way ANOVA | GC-P vs SGC-P p=0.0846 |
| | 1 Week sIPSC Amplitude (pA) | 17.35 ± 1.54 | 23.12 ± 1.12 | 21.27 ± 3.59 | 24.06 ± 1.54 | Interaction: F (1, 31) = 0.4471 P=0.5087 | **GC-S vs GC-P p=0.0643** |
| | | | | | | Cell Type: F (1, 31) = 1.176 P=0.2865 | SGC-S vs SGC-P p=0.4050 |
| | | | | | | **TreatmentF (1, 31) = 3.673 P=0.0646** | GC-S vs SGC-S p=0.2028 |
| | | | | | | by Two way ANOVA | GC-P vs SGC-P p=0.7802 |
| | 1 Week sIPSC Biexp fit Decay (ms) | 6.33 ± 0.36 | 6.07 ± 0.43 | 6.33 ± 0.48 | 5.41± 0.26 | Interaction: F (1, 31) = 0.6568 P=0.4239 | GC-S vs GC-P p =0.9850 |
| | | | | | | Cell Type: F (1, 31) = 0.6511 P=0.4259 | SGC-S vs SGC-P p=0.4533 |
| | | | | | | Treatment: F (1, 31) = 2.037 P=0.1635 | GC-S vs SGC-S p>0.9999 |
| | | | | | | by Two way ANOVA | GC-P vs SGC-P p=0.7346 |
| | 1 Week sIPSC Area Under the Curve (pA ms) | 79.33 ± 14.49 | 110.3 ± 8.95 | 111.8 ± 22.32 | 98.46 ±9.12 | Interaction: F (1, 31) = 2.040 P=0.1632 | GC-S vs GC-P p=0.4745 |
| | | | | | | Cell Type: F (1, 31) = 0.4448 P=0.5098 | SGC-S vs SGC-P p=0.9641 |
| | | | | | | Treatment: F (1, 31) = 0.3227 P=0.5741 | GC-S vs SGC-S p=0.4274 |
| | | | | | | by Two way ANOVA | GC-P vs SGC-P p=0.9769 |
| | 1 Week sIPSC Rise Time (ms) | 1.38 ± 0.08 | 1.21 ± 0.09 | 1.48 ± 0.1 | 1.06± 0.05 | Interaction: F (1, 31) = 2.273 P=0.1417 | GC-S vs GC-P p=0.5096 |
| | | | | | | Cell Type: F (1, 31) = 0.09037 P=0.7657 | **SGC-S vs SGC-P p=0.0087** |
| | | | | | | **Treatment: F (1, 31) = 11.78 P=0.0017** | GC-S vs SGC-S p=0.8496 |
| | | | | | | by Two way ANOVA | GC-P vs SGC-P p=0.6491 |

### Supplemental Table 5

| Figure | Parameter (Unit) | Control GC (GC-S) n=9/ 5 mice | Post-SE GC (GC-P) n=11/ 6 mice | Control SGC (SGC-S) n=9/ 5 mice | Post-SE SGC (SGC-P) n=10/ 6 mice | F & P value | Multiple Comparisons |
|---|---|---|---|---|---|---|---|
| 4 | 1 Month sIPSC Frequency (Hz) | 5.41± 1.5 | 7.15 ±0.84 | 6.9 ± 0.57 | 8.49 ±1.26 | Interaction:F (1, 36) = 0.004778 P=0.9453<br>Cell Type:F (1, 36) = 1.587 P=0.2158<br>Treatment:F (1, 36) = 2.204, P=0.1463<br>by Two way ANOVA | GC-S vs GC-P p=0.2670<br>SGC-S vs SGC-P p=0.3352<br>GC-S vs SGC-S p=0.3652<br>GC-P vs SGC-P p=0.3933 |
| | 1 Month sIPSC Amplitude (pA) | 16.12± 1.8 | 26.93 ±2.65 | 24.83 ±3.42 | 29.4 ± 1.17 | Interaction:F (1, 36) = 1.028 P=0.3175<br>**Cell Type:F (1, 36) = 3.299 P=0.0777**<br>**Treatment: F (1, 36) = 6.244, P=0.0172**<br>by Two way ANOVA | **GC-S vs GC-P p=0.0152**<br>SGC-S vs SGC-P p=0.3124<br>**GC-S vs SGC-S p=0.0587**<br>GC-P vs SGC-P p=0.5640 |
| | 1 Month sIPSC Biexp fit Decay (ms) | 7.18 ± 0.46 | 6.81 ±0.45 | 5.699 ± 0.46 | 5.94 ± 0.41 | Interaction: F (1, 36) = 0.4614 P=0.5013<br>**Cell Type: F (1, 36) = 6.874 P=0.0127**<br>Treatment: F (1, 36) = 0.02046 P=0.8871<br>by Two way ANOVA | GC-S vs GC-P p=0.9606<br>SGC-S vs SGC-P p=0.9933<br>GC-S vs SGC-S p=0.1103<br>GC-P vs SGC-P p=0.5191 |
| | 1 Month sIPSC Area Under the Curve (pA ms) | 54.49 ±11.62 | 110.8 ±17.87 | 110.6 ±14.32 | 115.1 ±16.10 | Interaction: F (1, 36) = 2.806 P=0.1026<br>Cell Type: F (1, 36) = 3.833 P=0.0580<br>Treatment: F (1, 36) = 3.881 P=0.0566<br>by Two way ANOVA | **GC-S vs GC-P p=0.0473**<br>SGC-S vs SGC-P p=0.9993<br>GC-S vs SGC-S p=0.0657<br>GC-P vs SGC-P p=0.9993 |
| | 1 Month sIPSC Rise Time (ms) | 1.28 ±0.11 | 1.33 ±0.12 | 1.64 ±0.15 | 1.12 ±0.06 | **Interaction: F (1, 36) = 6.341 P=0.0164**<br>Cell Type: F (1, 36) = 0.4792 P=0.4932<br>**Treatment: F (1, 36) = 4.417 P=0.0426**<br>by Two way ANOVA | GC-S vs GC-P p=0.9969<br>**SGC-S vs SGC-P p=0.0118**<br>GC-S vs SGC-S p=0.1262<br>GC-P vs SGC-P p=0.5771 |

### Supplemental Table 6

| Figure | | Stimulus Amplitude | Control GC | Post-SE GC | Control SGC | Post-SE | F & P value |
|---|---|---|---|---|---|---|---|
| 5 | 1 Week eEPSCs Amplitude | 0.5 | 133.07±41.96 | 157.54 ± 84.95 | 171.82 ± 76.59 | 131.65±69.08 | Stim Intensity: F (3, 120) = 39.45 P<0.0001<br>Treatment: F (1, 120) = 2.082 P=0.1517 |
| | | 1 | 479.58±104.42 | 501.37±135.90 | 513.09±119.43 | 542.78±183.02 | Cell Type: F (1, 120) = 0.01193 P=0.9132<br>Stim Intensity x Treatment: F (3, 120) = 0.6569 P=0.5802 |
| | | 1.5 | 831.49± 134.90 | 1027.32±172.09 | 826.81± 108.78 | 946.65±223.32 | Stim Intensity x Cell Type: F (3, 120) = 0.06297 P=0.9793<br>Treatment x Cell Type: F (1, 120) = 0.05021 P=0.8231 |
| | | 2 | 1103.95± 148.3 | 1354.61±220.43 | 1070.33±94.00 | 1321.23±263.65 | Stim Intensity x Treatment x Cell Type: F (3, 120) = 0.02146 P=0.9957 by 3Way ANOVA |
| | 1 Month eEPSCs Amplitude | 0.5 | 83.76 ± 28.11 | 90.04± 31.12 | 204.22± 68.85 | 181.17± 59.43 | Stim Intensity: F (3, 128) = 48.87 P<0.0001<br>Treatment: F (1, 128) = 7.084 P=0.0088 |
| | | 1 | 485.59 ± 128.18 | 380.47± 81.20 | 556.43± 165.6 | 485.46± 86.99 | Cell Type F (1, 128) = 3.105 P=0.0805<br>Stim Intensity x Treatment: F (3, 128) = 1.376 p=0.2530 |
| | | 1.5 | 853.78 ± 195.76 | 698.76± 112.68 | 1019.52± 146.27 | 732.61± 75.29 | Stim Intensity x Cell Type: F (3, 128) = 0.02872 P=0.9934<br>Treatment x Cell Type: F (1, 128) = 0.9895 P=0.3217 |
| | | 2 | 1132.07± 223.84 | 979.22± 116.60 | 1447.69± 167.23 | 936.46± 117.90 | Stim Intensity x Treatment x Cell Type: F (3, 128) = 0.4980 P=0.6843 by 3Way ANOVA |

### Supplemental Table 7

| Figure | | Stimulus Amplitude | Control GC | Post-SE GC | Control SGC | Post-SE | F & P value |
|---|---|---|---|---|---|---|---|
| 5 | 1 Week eIPSCs Amplitude | 0.5 | 57.02 ± 21.21 | 122.52 ± 38.76 | 274.69 ± 98.96 | 114.24 ± 54.93 | Stim Intensity: F (3, 120) = 39.44 P<0.0001<br>Treatment: F (1, 120) = 8.436 P=0.0044 |
| | | 1 | 271.22 ± 81.90 | 317.08 ± 89.07 | 608.31 ± 121.43 | 251.88 ± 81.97 | Cell Type: F (1, 120) = 11.78 P=0.0008<br>Stim Intensity x Treatment: F (3, 120) = 0.5156 p=0.6723 |
| | | 1.5 | 606.49 ± 87.11 | 560.50 ± 145.00 | 972.69 ± 126.57 | 618.59 ± 95.05 | Stim Intensity x Cell Type: F (3, 120) = 0.5545 P=0.6461<br>Treatment x Cell Type: F (1, 120) = 8.336 P=0.0046 |
| | | 2 | 791.45 ± 82.202 | 722.37 ± 171.75 | 1224.37 ± 163.56 | 854.42 ± 66.37 | Stim Intensity x Treatment x Cell Type: F (3, 120) = 0.1138 P=0.9519 by 3Way ANOVA |
| | 1 Month eIPSCs Amplitude | 0.5 | 51.78 ± 15.30 | 63.62 ± 13.27 | 129.86 ± 87.46 | 112.76 ± 42.78 | Stim Intensity: F (3, 128) = 46.50 P<0.0001<br>Treatment: F (1, 128) = 0.3409 P=0.5604 |
| | | 1 | 364.46 ± 87.45 | 364.95 ± 109.48 | 424.57 ± 112.44 | 432.21 ± 96.83 | Cell Type: F (1, 128) = 3.123 P=0.0796<br>Stim Intensity x Treatment: F (3, 128) = 0.1427 P=0.9342 |
| | | 1.5 | 655.88 ± 105.45 | 637.14 ± 126.15 | 801.11 ± 134.44 | 749.63 ± 128.19 | Stim Intensity x Cell Type: F (3, 128) = 0.1323 P=0.9407<br>Treatment x Cell Type: F (1, 128) = 0.3712 P=0.5434 |
| | | 2 | 860.28 ± 135.50 | 872.17 ± 145.45 | 1089.76 ± 143.98 | 902.20 ± 118.21 | Stim Intensity x Treatment x Cell Type: F (3, 128) = 0.1747 P=0.9134 by 3Way ANOVA |
