## Supplemental Figure for "Altered Physiology and Ensemble Recruitment of Dentate Gyrus Semilunar Granule Cells in a Mouse Model of Epilepsy"

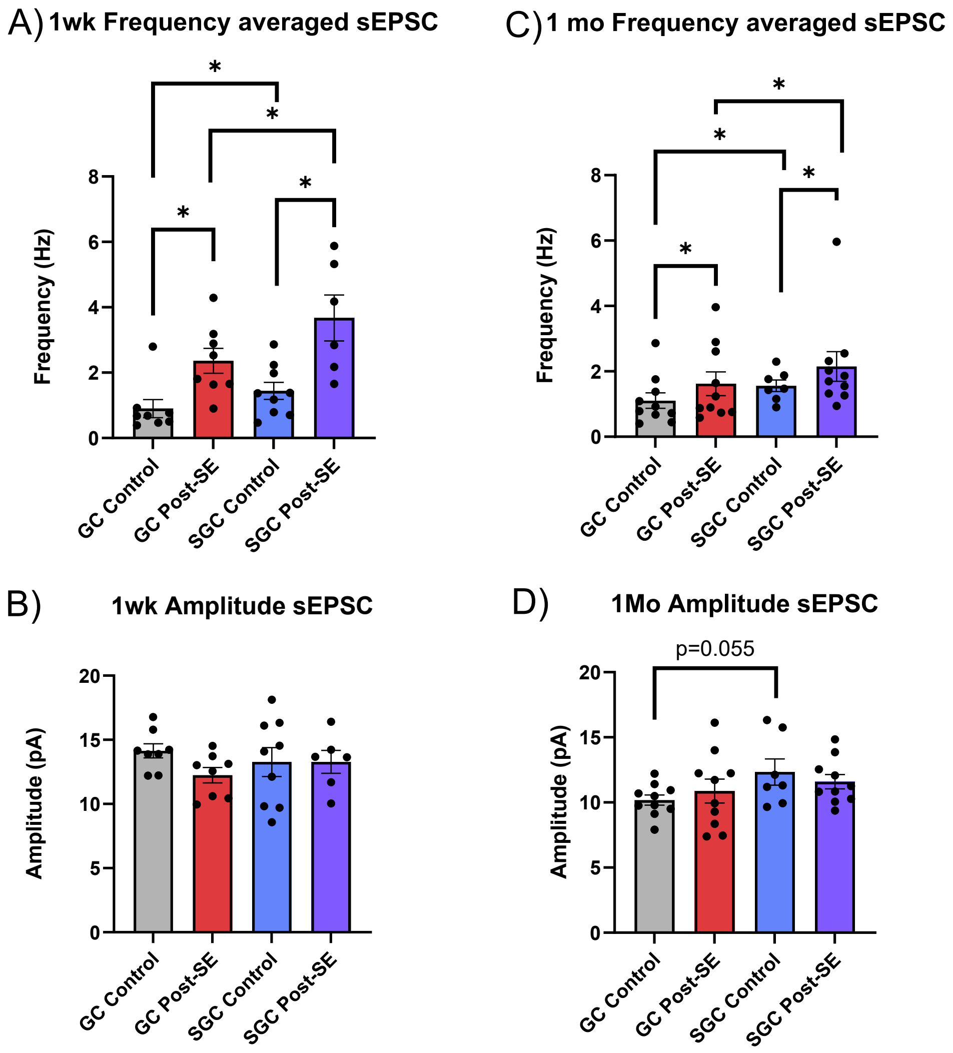


**Supplemental Figure 1: Spontaneous EPSC frequency in both GCs and SGCs is increased after SE.** A-B) Summary plots of cell-averaged sEPSC frequency (A) and amplitude (B) in GCs and SGCs 1-week after SE and in age-matched controls. C-D) Cell-averaged sEPSC frequency (C) and amplitude (D) in GCs and SGCs 1-month after SE and in age-matched controls. *indicates $p<0.05$ TW-ANOVA with Šídák's multiple comparisons post hoc tests.

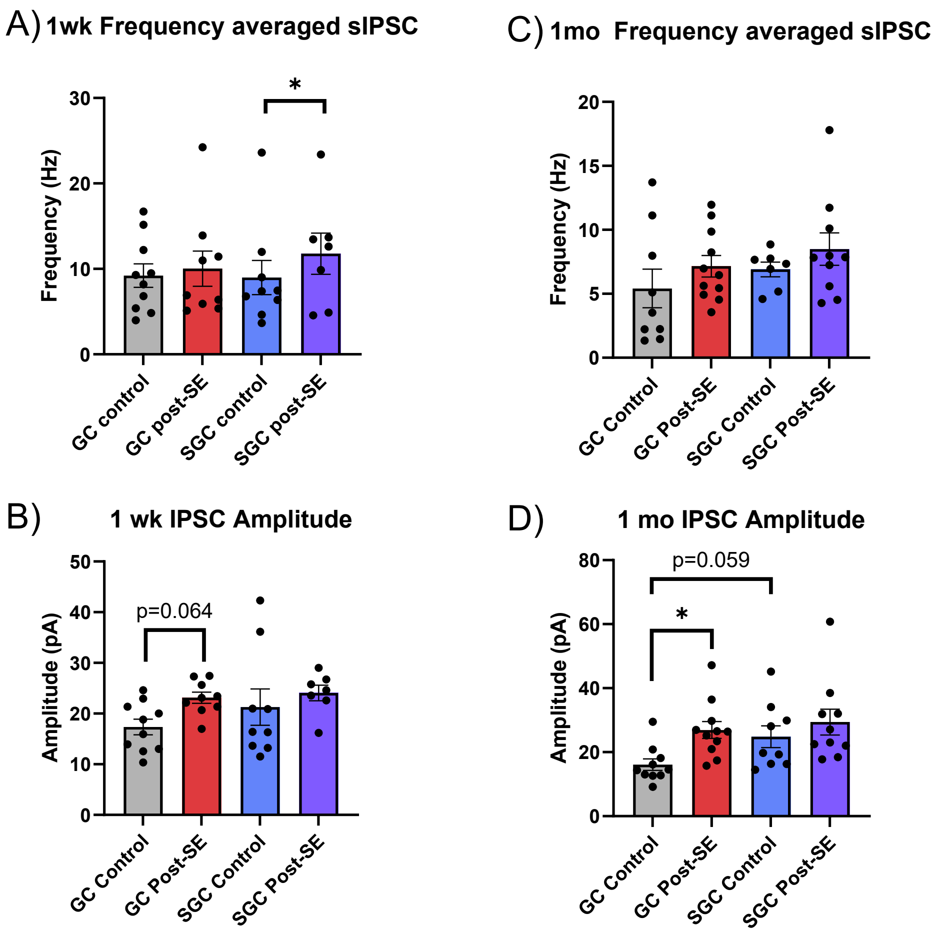


**Supplemental Figure 2: Spontaneous IPSC frequency in GCs and SGCs after SE.** A-B) Summary plots of cell-averaged sIPSC frequency (A) and amplitude (B) in GCs and SGCs 1-week after SE and in age-matched controls. C-D) Cell-averaged sIPSC frequency (C) and amplitude (D) in GCs and SGCs 1-month after SE and in age-matched controls. *indicates $p<0.05$ TW-ANOVA with Šídák's multiple comparisons post hoc tests.
